# An Open Test–Retest Dataset of Rapid Myelin Mapping Acquisitions

**DOI:** 10.64898/2026.09.29.753255

**Authors:** Steven L. Meisler, Taylor Salo, Matthew Cieslak, Brooke L. Sevchik, Juliette Brook, Guillaume Duhamel, Olivier M. Girard, Bart Larsen, Shraddha Pandey, Lucinda M. Sisk, Lucas Soustelle, Valerie J. Sydnor, Poljanka Johnson, David R. Roalf, Russell T. Shinohara, Manuel Taso, M. Dylan Tisdall, Theodore D. Satterthwaite

**Author notes:** Correspondence to Theodore D. Satterthwaite. Co-first authors who contributed equally (Steven L. Meisler, Taylor Salo). Co-senior authors who contributed equally (Manuel Taso, M. Dylan Tisdall, Theodore D. Satterthwaite).

## Abstract

Myelin supports rapid and efficient neural communication and is implicated in brain development, learning, aging, and neurological disease. Although numerous magnetic resonance imaging (MRI) approaches have been proposed to characterize myelin in vivo, they have rarely been evaluated together within the same participants and across repeated acquisitions. To address these gaps, we introduce the Myelin Imaging Reliability and ReprOducibility Resource (MIRROR), an openly available multimodal test– retest MRI dataset of diffusion MRI (dMRI), inhomogeneous magnetization transfer imaging (ihMT), quantitative susceptibility mapping (QSM), MP2RAGE, T_1_w/T_2_w ratio mapping, and transverse relaxometry in 22 healthy adults. We evaluated gray–white matter differentiation, intermetric correspondence, test–retest reliability, and participant specificity across 27 myelin-sensitive metrics. Gray–white matter differentiation was strong for dMRI derivatives, MP2RAGE-derived R_1_, T_1_w/T_2_w, and ihMT, while being substantially weaker for QSM measures. Intermetric spatial correlations were generally strongest within acquisition or modeling families. Test–retest reliability was generally high, particularly for dMRI-derived metrics, but was more variable among QSM-, ihMT-, and R_2_*-derived measures. Regional spatial profiles were also highly participant-specific, with discriminability approaching its upper bound for most metrics. All imaging data, derived maps, and processing code are openly released. MIRROR provides a benchmark for characterizing the measurement properties and practical tradeoffs of scalable myelin-sensitive MRI methods, supporting principled acquisition selection and accelerating methodological development and validation.

## 1. Introduction

Myelin is essential for rapid, efficient, and synchronous neural communication (Dubey et al., 2022; Pajevic et al., 2014). Although myelination was historically thought to stabilize after early development, accumulating evidence suggests that myelin remains dynamic across the lifespan and may support learning, memory, and adaptation (Bacmeister et al., 2022; Eg et al., 2018; Fields, 2008, 2015; Pan et al., 2020). In humans, white matter continues to mature into early adulthood before declining later in life (Chapman & Hill, 2020). Loss of myelination is also a defining feature of neurological diseases such as multiple sclerosis (Steinman, 1996). Patterns of myelination have been associated with large-scale cortical organization (Huntenburg et al., 2017; Paquola et al., 2019), cognitive traits (NSPN Consortium et al., 2019), psychiatric risk (Nave & Ehrenreich, 2014; Ziegler et al., 2019), and experience-dependent plasticity (Bonetto et al., 2021; Fields, 2015; Ramos-Cabrer et al., 2025). Characterizing myelin in vivo therefore offers a powerful opportunity to study brain health across the lifespan, with many applications to cognitive and clinical neuroscience. However, despite the growing interest in myelin mapping, the proliferation of available measures has outpaced systematic benchmarking of their relative sensitivity, specificity, and test–retest reliability. Here we introduce a new open test–retest data resource for multiple myelin-sensitive MRI modalities.

Signal from many of the macromolecular protons abundant in myelin decays on sub-millisecond timescales, making it largely unresolvable to conventional MRI sequences (Piredda et al., 2021). Specialized approaches such as ultrashort echo time MRI (Du et al., 2014) may detect short T_2_ myelin-associated signal, but many commonly used myelin-sensitive measures rely on indirect contrasts that may reflect distinct biological and physical properties. Magnetization transfer methods probe exchange between free-water protons and protons bound to macromolecules, including the lipids and proteins abundant in myelin (Alsop et al., 2023; Duhamel et al., 2019; Wolff & Balaban, 1989). Quantitative T_1_ relaxometry is sensitive to dipolar interactions between water and tissue macromolecules, including those associated with myelin, although iron and water content also influence longitudinal relaxation (Lutti et al., 2014; Stüber et al., 2014). T_1_w/T_2_w ratios are similarly sensitive to nonspecific myelin-related differences in tissue relaxation, with greater myelination tending to increase T_1_-weighted signal and decrease T_2_-weighted signal (Baum et al., 2022; Glasser & Van Essen, 2011). Multi-echo T_2_ relaxometry can estimate short-T_2_ water components associated with water confined between myelin bilayers (Alonso-Ortiz et al., 2015). Diffusion MRI is indirectly sensitive to myelin because myelin sheaths and other cellular membranes restrict and shape water diffusion within tissue (Laule et al., 2007). Finally, quantitative susceptibility mapping is sensitive to myelin because myelin-forming lipids are diamagnetic and contribute orientation-dependent magnetic susceptibility, although susceptibility is also strongly influenced by paramagnetic iron (Liu et al., 2015). Consequently, different myelin-sensitive MRI measures often show only partial agreement, complicating biological interpretation and limiting the synthesis of findings across studies (Lazari & Lipp, 2021; Mancini et al., 2020; van der Weijden et al., 2021).

Existing myelin neuroimaging studies are often limited to a single or small set of imaging modalities and a single acquisition per participant. Moreover, many myelin-sensitive contrasts lack the mature and standardized processing ecosystems available for widely used diffusion analyses (Cieslak et al., 2021). These limitations impede efforts to compare the reliability, convergence, and tissue-specific behavior of existing myelin mapping methods. There is therefore a need for openly available, fully processed multimodal datasets that enable complementary myelin-sensitive measures to be evaluated throughout the brain across repeated acquisitions.

To address these needs, we present the MIRROR (Myelin Imaging Reliability and ReprOducibility Resource) dataset, an openly available multimodal MRI resource for benchmarking in vivo myelin-sensitive neuroimaging methods. MIRROR comprises several complementary acquisitions that can be incorporated into conventional neuroimaging sessions, reflecting the need for scalable protocols in developmental, clinical, and population studies where scan time may be limited. The protocol includes diffusion MRI, inhomogeneous magnetization transfer imaging (ihMT), quantitative susceptibility mapping (QSM), quantitative R_1_ estimation, T_1_w/T_2_w ratio mapping, and transverse relaxometry, with each acquisition repeated to enable test–retest evaluation. Using these data, we assess the degree of gray–white matter differentiation, intermetric correspondence, test– retest reliability, and participant specificity across 27 putative myelin-sensitive measures. We also openly release the imaging data, derived maps, and all processing code to facilitate reproducible methodological development and benchmarking. Together, MIRROR provides a framework for characterizing myelin-related variation and evaluating scalable approaches to in vivo myelin imaging.

## 2. Methods

### 2.1 Participants and Study Design

Twenty-two adults participated in the primary study sample (15 female; mean age = 23.95 years, SD = 3.77; range = 20–30 years). All study protocols were approved by the University of Pennsylvania Institutional Review Board (Protocol 857664 / 24-2318). All participants provided written informed consent to enroll in the study. Exclusion criteria included a history of head trauma with significant loss of consciousness; neurological or medical conditions that could affect brain anatomy, physiology, or cognitive performance; pregnancy; insufficient English proficiency to understand consent or study instructions; MRI contraindications; and recent substance use associated with possible acute intoxication or withdrawal. Each participant completed two identical MRI sessions on the same day. After the first session, participants exited the scanner for no more than 10 minutes and were then repositioned for the second session. An additional four pilot participants completed a single MRI session; these data are shared but were not included in analyses.

### 2.2 Data Acquisition

The complete MRI protocol is summarized in **Table 1**. The source acquisitions required to generate each derivative family are summarized in **Supplementary Figure S1**, with their cumulative acquisition durations reported in **Supplementary Table S1**. The protocol included conventional anatomical sequences, diffusion-weighted MRI (dMRI), and several specialized acquisitions from which myelin-sensitive measures were derived. All data were collected on a single Siemens Healthineers Prisma 3T scanner using a 64-channel head and neck coil. Data were organized according to the Brain Imaging Data Structure (BIDS; K. J. Gorgolewski et al., 2016) wherever an applicable specification was available. Because ihMT is not yet represented in an official BIDS extension, these data were organized using analogous BIDS-like naming and directory conventions.

**Table 1.** MRI acquisition parameters for each session. Multiple entries within a cell indicate that an acquisition comprised multiple scans.

| Acquisition | Acquisition Time (MM:SS) | Echo Time (ms) | Repetition Time (s) | Field-of-View (mm) | Voxel Size (mm) |
| --- | --- | --- | --- | --- | --- |
| MPRAGE T<sub>1w</sub> | 4:05 | 2.81 | 2.3 | 256×256 | 1×1×1 |
| SPACE T<sub>1w</sub> | 4:11 | 9 | 0.7 | 256×256 | 1×1×1 |
| SPACE T<sub>2w</sub> | 4:04 | 133 | 2 | 256×256 | 1×1×1 |
| B<sub>1</sub><sup>+</sup> field map | 0:21 | 2.06 | 9.5 | 256×256 | 4×4×4 |
| MP2RAGE | 7:59 | 2.81 | 4.5 | 256×256 | 1×1×1 |
| ihMTRAGE | 8:22 | 2.9 | 2 | 256×176 | 2×2×2 |
| dMRI | 12:48 | 88 | 4.8 | 238×238 | 1.7×1.7×1.7 |
| AP | 6:24 |  |  |  |  |
| PA | 6:24 |  |  |  |  |
| MESE | 3:22 |  |  | 230×230 | 0.9×0.9×3 |
| AP TE1 | 0:40 | 15 | 5.71 |  |  |
| AP TE2 | 0:40 | 30 | 5.71 |  |  |
| AP TE3 | 0:40 | 50 | 5.71 |  |  |
| AP TE4 | 0:42 | 100 | 6.03 |  |  |
| PA TE1 | 0:40 | 15 | 5.71 |  |  |
| MEGRE | 5:37 | 6.34, 11.78,<br>17.32, 22.81,<br>28.30 | 0.033 | 256×256 | 1×1×1 |

#### 2.2.1. T1-weighted MPRAGE

One T_1_w gradient echo (3D MPRAGE) scan was acquired each session with the following parameters: isotropic spatial resolution of 1 mm; echo time (TE) of 2.81 ms; repetition time (TR) of 2.3 s; flip angle of 8°; field of view (FOV) of 256×256 mm; 176 slices; total acquisition time of 4:05. 3D distortion correction and prescan normalization filters were applied by the scanner. These data were named according to the following convention:

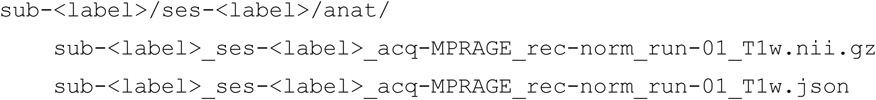

#### 2.2.2. T1-weighted SPACE

One T_1_w 3D SPACE scan was acquired each session with the following parameters: isotropic spatial resolution of 1 mm; TE of 9 ms; TR of 0.7 s; flip angle of 100°; FOV of 256×256 mm; 176 slices; total acquisition time of 4:11. The raw, 3D distortion correction, and prescan normalization filters were all applied by the scanner. These data were named according to the following convention:

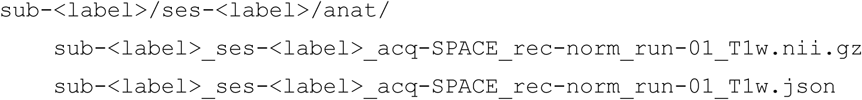

#### 2.2.3. T2-weighted SPACE

One T_2_w 3D SPACE scan was acquired each session with the following parameters: isotropic spatial resolution of 1 mm; TE of 133 ms; TR of 2 s; flip angle of 100°; FOV of 256×256 mm; 176 slices; total acquisition time of 4:04. The raw, 3D distortion correction, and prescan normalization filters were all applied by the scanner. These data were named according to the following convention:

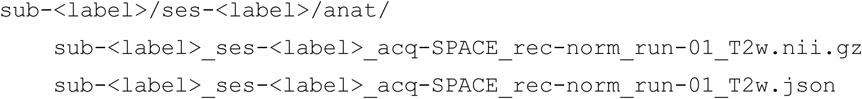

#### 2.2.4. B1^+^ field map

One Siemens product presaturated turbo-flash (TB1TFL) scan (Chung et al., 2010) was acquired each session to measure the B_1_ transmit (B_1_^+^) field, with the following parameters: isotropic spatial resolution of 4 mm; TE of 2.06 ms; TR of 9.5 s; flip angle of 8°; FOV of 256×256 mm; 34 slices; total acquisition time of 0:21. No filters were applied at the scanner. These data were named according to the following convention:

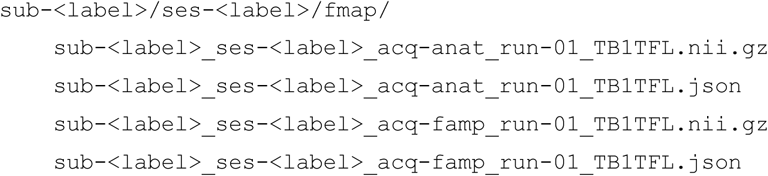

#### 2.2.5. MP2RAGE

One 3D MP2RAGE (Marques et al., 2010) scan was acquired each session with the following parameters: isotropic spatial resolution of 1 mm; TE of 2.81 ms; TR of 4.5 s; inversion times (TIs) of 0.7 and 2.2 s; flip angles of 4° and 5°; FOV of 256×256 mm; 176 slices; total acquisition time of 7:59. The raw, 3D distortion correction, and prescan normalization filters were applied by the scanner. Magnitude and uniform T_1_-weighted (UNIT1) images were available for all participants. Phase images were reconstructed for both sessions in 8 participants and one session in 2 participants. These data were named according to the following convention:

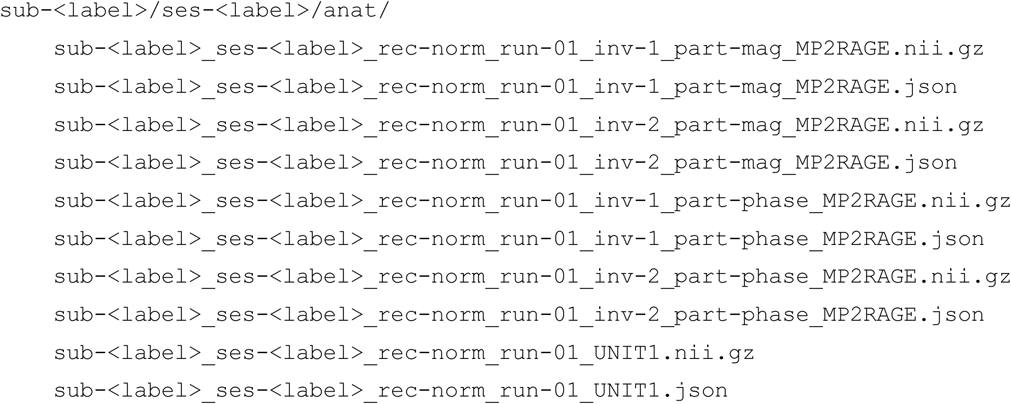

#### 2.2.6. Inhomogeneous Magnetization Transfer with RApid Gradient Echo

One 3D inhomogeneous Magnetization Transfer with RApid Gradient Echo (ihMTRAGE; Varma et al., 2020) scan was acquired each session with an isotropic spatial resolution of 2 mm, TE of 2.9 ms, TR of 2 s, flip angle of 7°, FOV of 256 × 176.128 mm, 80 slices, and a total acquisition time of 8:22. The acquisition comprised an unsaturated reference (M_0_) volume, single-frequency saturation at positive and negative offsets, and two dual-frequency saturation volumes. Each ihMT preparation consisted of ten 5-ms Tukey-shaped saturation pulses with a nominal flip angle of 975°, delivered one pulse per burst with a burst repetition time of 100 ms, for a total preparation duration of 910 ms. Single-frequency saturation was applied at offsets of ±7 kHz, whereas dual-frequency saturation used cosine-modulated RF pulses to simultaneously apply symmetric saturation at ±7 kHz. The raw and 3D distortion correction filters were all applied by the scanner. These data were named according to the following convention:

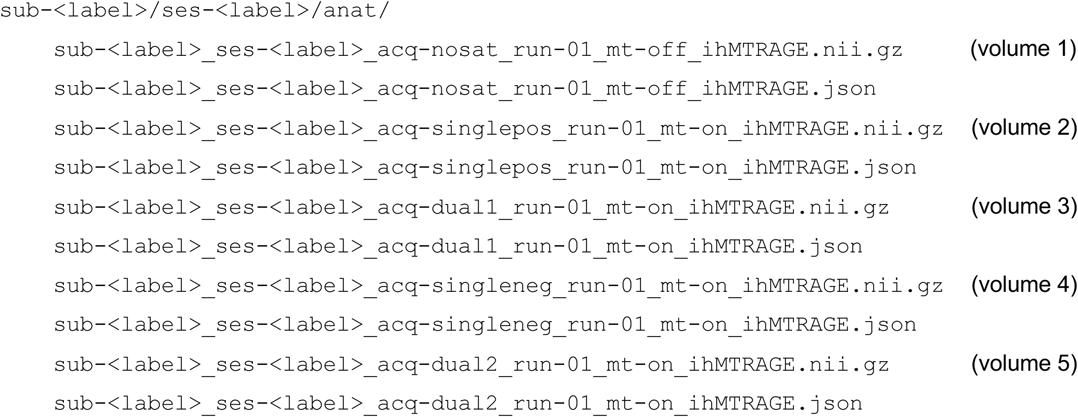

#### 2.2.7. Diffusion MRI (dMRI)

Two runs of dMRI data were acquired each session following the HEALthy Brain and Child Development (HBCD) protocol (Cieslak et al., 2025; Dean et al., 2024). The HBCD dMRI protocol includes two acquisitions with opposite phase-encoding directions (AP and PA) and the following parameters: isotropic spatial resolution of 1.7 mm; TE of 88 ms; TR of 4.8 s; flip angle of 78°; FOV of 238×238 mm; 87 slices; *b*-values of 0 (11 volumes), 500 (6 directions), 1000 (12 directions), 2000 (18 directions), and 3000 (29 directions) s/mm^2^; total acquisition time of 12:48 across both scans. Prescan normalization was applied to the magnitude data by the scanner. These data were named according to the following convention:

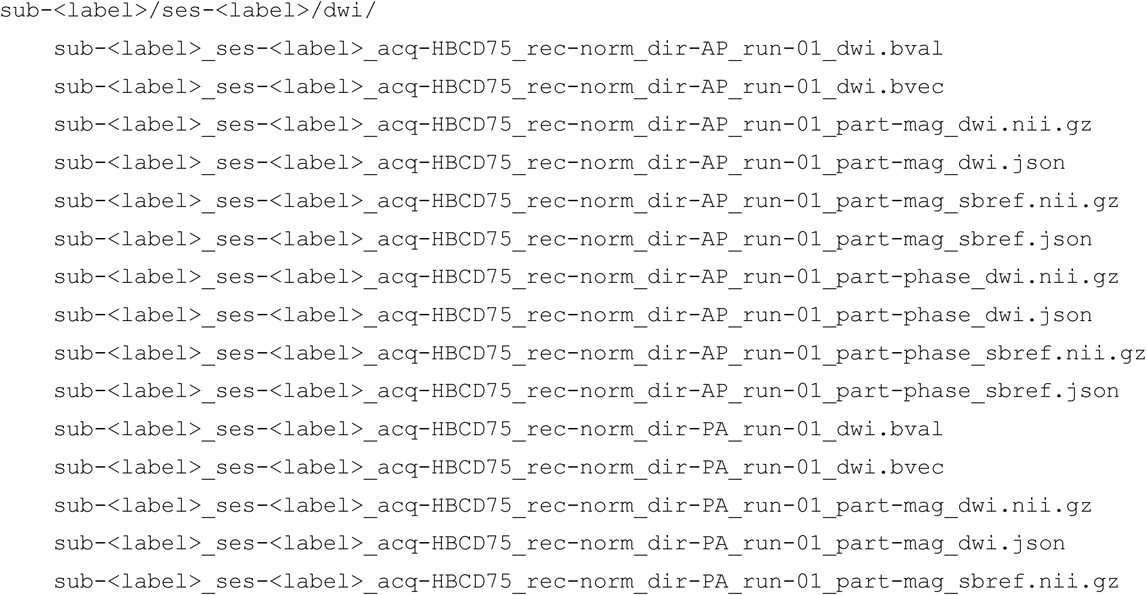

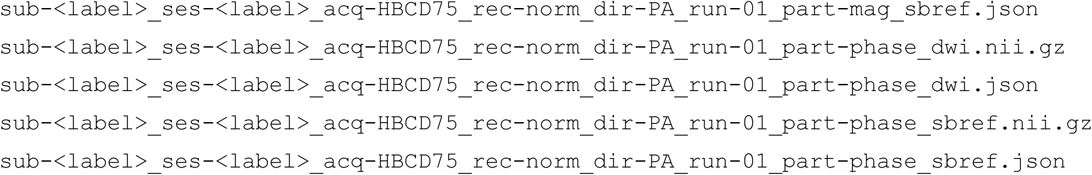

#### 2.2.8. Multi-echo Spin Echo (MESE)

Four 2D segmented spin echo EPI scans were acquired each session with different TEs, so that the four volumes could be combined and used as a multi-echo spin-echo acquisition. Each spin echo scan had slightly different acquisition parameters: the first three spin echo scans were acquired with the following parameters: voxel size of 0.9×0.9×3 mm; TR of 5.71 s; flip angle of 90°; FOV of 230×230 mm; 48 slices; acquisition time of 0:40. The TEs were 15 ms, 30 ms, and 50 ms. The fourth spin echo scan was acquired with the following parameters: voxel size of 0.9×0.9×3 mm; TR of 6.03 s; flip angle of 90°; FOV of 230×230 mm; 48 slices; total acquisition time of 0:42; and TE of 100 ms.

One additional scan was acquired each session with a reversed phase encoding direction (PA), otherwise matching the settings of the spin echo scan with a TE of 15 ms, for susceptibility-distortion correction. Due to acquisition errors, four participants received an AP acquisition instead of the intended PA acquisition during one session. Because valid reverse-phase-encoded data were therefore unavailable for all participants and sessions, PA images were not incorporated into the primary processing pipeline, although all acquired images are released. The total acquisition time across all five scans was 3:22 per session. No filters were applied at the scanner. These data were named according to the following convention:

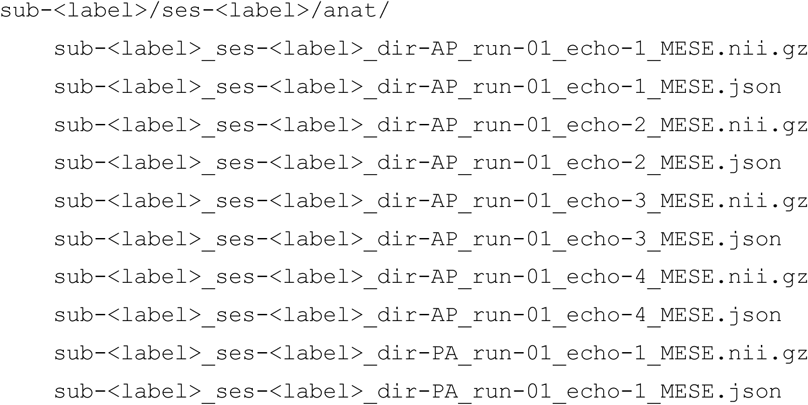

#### 2.2.9. Multi-echo gradient-recalled echo (MEGRE)

One 3D MEGRE scan was acquired each session with the following parameters: isotropic spatial resolution of 1 mm; TEs of 6.34, 11.78, 17.32, 22.81, and 28.30 ms; TR of 0.033 s; flip angle of 15°; FOV of 256×256 mm; magnitude and phase image reconstruction; 144 slices; total acquisition time of 5:37. Due to an error in the sequence programming, the first echo of the MEGRE scan had a pixel bandwidth of 310 Hz/pixel, while the other echoes had bandwidths of 250 Hz/pixel. The 3D distortion correction filter was applied by the scanner. These data were named according to the following convention:

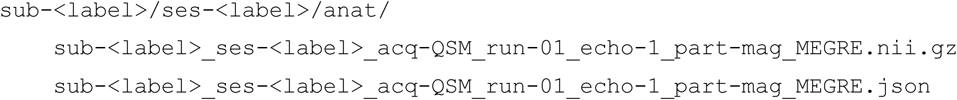

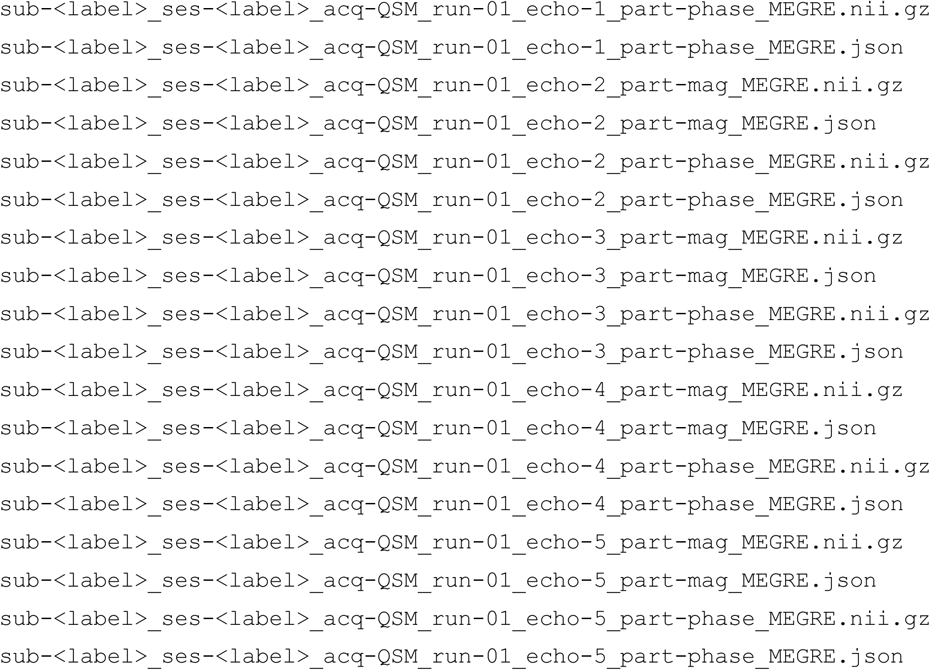

### 2.3. Data Anonymization

Prior to processing, MPRAGE T_1_w and SPACE T_1_w images were refaced with AFNI’s @afni_refacer_run (R. Cox & Taylor, 2020; R. W. Cox, 1996; R. W. Cox & Hyde, 1997). SPACE T_2_w and MP2RAGE images were defaced with pydeface (Gulban et al., 2022). The other modalities lacked sufficient resolution, contrast, or field of view to necessitate defacing. We additionally removed image metadata that might allow reidentification, such as the acquisition dates and times for the sessions.

### 2.4. Data Processing

The data were processed with a combination of standardized tools. Additionally, custom Python code was used to coordinate processing scripts, handle intermodal coregistration to the fMRIPrep-preprocessed T_1_w image (henceforth referred to as “T_1_w space”), and transform derivatives into common spaces for analysis. Throughout all analyses, continuous-valued derivative metric maps were transformed to T_1_w and ICBM 152 nonlinear asymmetric template spaces (version 2009c, TemplateFlow ID: MNI152NLin2009cAsym, henceforth referred to as “MNI space”) (Fonov et al., 2009) using linear interpolation.

#### 2.4.1. MPRAGE Processing

Preprocessed MPRAGE T_1_w images were used to inform coregistrations and spatial normalization. The MPRAGE T_1_w images were first minimally preprocessed using the anatomical processing workflow in fMRIPrep version 25.0.0 (Esteban et al., 2019; Markiewicz et al., 2025), which is based on Nipype 1.9.2 (Esteban et al., 2025; K. Gorgolewski et al., 2011). This workflow produced a participant-averaged preprocessed MPRAGE T_1_w, as well as T_1_w-to-MNI transforms for each participant. The following paragraphs contain partially edited text generated by fMRIPrep, which is distributed with a Creative Commons Zero (CC0) license with the expressed intention of being included in manuscripts.

Each T_1_w image was corrected for intensity non-uniformity (INU) with N4BiasFieldCorrection (Tustison et al., 2010), distributed with ANTs 2.5.4 (Avants et al., 2008). For participants with multiple T_1_w images, these INU-corrected images were combined to an unbiased participant average using FreeSurfer’s mri_robust_template functionality. Images were then skull-stripped with a Nipype implementation of the antsBrainExtraction.sh workflow (from ANTs), using OASIS30ANTs as the target template. Brain tissue segmentation of cerebrospinal fluid (CSF), white matter (WM) and gray matter (GM) was performed on the brain-extracted T_1_w using fast from FSL version 6.0.7.13 (Y. Zhang et al., 2001). Brain surfaces were reconstructed using recon-all from FreeSurfer version 7.3.2 (Dale et al., 1999). The brain mask estimated previously was refined with a custom variation of the method used by Mindboggle (Klein et al., 2017) to reconcile ANTs-derived and FreeSurfer-derived segmentations of cortical gray matter. Volume-based spatial normalization to MNI template space was performed through nonlinear registration with antsRegistration, using brain-extracted versions of both the participant and template T_1_w images.

#### 2.4.2. T1w/T2w Ratio Processing

The raw MPRAGE T_1_w, SPACE T_1_w, and SPACE T_2_w images were used to derive scaled and unscaled T_1_w/T_2_w ratio maps. All three images were first skullstripped with SynthStrip version 1.7 and coregistered to the preprocessed MPRAGE T_1_w using rigid-body alignment with ANTs. The unscaled versions of the T_1_w/T_2_w ratio maps were calculated by dividing the coregistered images. This resulted in two T_1_w/T_2_w ratio maps: unscaled MPRAGE T_1_w divided by unscaled SPACE T_2_w and unscaled SPACE T_1_w divided by unscaled SPACE T_2_w, hereafter referred to as “unscaled MPRAGE T_1_w/T_2_w” and “unscaled SPACE T_1_w/T_2_w”, respectively.

For the scaled versions of the T_1_w/T_2_w ratio maps, each raw image was scaled between 0 and 100 (Newman et al., 2024). To do this, each raw image was rescaled so that the minimum value across the whole image was set to 0 and the maximum value across the whole image was set to 100. The transforms estimated from the unscaled images were applied to the corresponding scaled images, after which the scaled ratios were calculated in T1w space. This resulted in two T1w/T_2_w ratio maps: scaled MPRAGE T_1_w divided by scaled SPACE T_2_w and scaled SPACE T_1_w divided by scaled SPACE T_2_w, hereafter referred to as “scaled MPRAGE T_1_w/T_2_w” and “scaled SPACE T_1_w/T_2_w”, respectively. The four T_1_w/T_2_w ratio maps were then transformed to T_1_w and MNI spaces.

#### 2.4.3. B1^+^ Field Map Processing

The TB1TFL sequence was used to measure the B_1_^+^ field map, which was subsequently used to correct MP2RAGE and ihMTRAGE outputs for transmit field inhomogeneity. The TB1TFL field map produced two images: an anatomically-weighted image and a scaled flip angle image. The anatomically-weighted image was used to register the field map to other images, while the scaled flip angle map was used to measure the difference between the expected flip angle and the real flip angle acquired by the scanner. The flip angle image was first divided by 10 times the nominal flip angle (79.99-80°) to express the measured transmit field relative to the prescribed flip angle. The anatomically-weighted image was then registered to the first inversion time magnitude image from the MP2RAGE scan using rigid-body alignment and the transform was retained.

#### 2.4.4. MP2RAGE Processing

The MP2RAGE scan was used to calculate quantitative T_1_ and R_1_ maps, both with and without B_1_^+^ bias field correction. UNIT1 and quantitative T_1_ maps were estimated from the MP2RAGE scan using a modified version of the pymp2rage Python library. Processing steps included masking of background voxels and conversion of UNIT1 image signal intensities to quantitative T1 values using sequence-specific lookup tables implemented in pymp2rage (Marques et al., 2010). We modified pymp2rage to use the signed UNIT1 image produced by the scanner instead of calculating a new one from complex-valued data. The quantitative T_1_ map was rescaled from milliseconds to seconds and then converted to a quantitative R_1_ map in Hertz. B_1_-corrected versions of the T_1_-weighted, quantitative T_1_, and quantitative R1 maps were also generated. The B_1_-corrected T1-weighted map was registered to T_1_w space, and the transform was retained in order to warp the quantitative R_1_ maps to T_1_w and MNI spaces.

#### 2.4.5. ihMTRAGE Processing

The ihMTRAGE scan was used to calculate several derivatives, including magnetization transfer ratio (MTR), inhomogeneous magnetization transfer image (ihMTw), inhomogeneous magnetization transfer ratio (ihMTR), inhomogeneous magnetization transfer saturation (ihMTsat), and inhomogeneous magnetization transfer saturation normalized by squared B_1_^+^ (ihMTsat-B_1_c). MTR provides a semi-quantitative measure of the overall motion-restricted macromolecular content. ihMTw is the ihMT signal selective to dipolar order generated in such environments, such as those found in myelin-rich tissue. ihMTR provides a semi-quantitative measure of the macromolecular content associated with dipolar order which has been shown to be more selective to myelin (Duhamel et al., 2019; Hertanu et al., 2022). ihMTsat captures the same underlying ihMT contrast using an effective metric designed to reduce T_1_ and signal-intensity biases. ihMTsat-B1c further normalizes ihMTsat by the squared transmit field to reduce bias from B_1_^+^ inhomogeneity. ihMTR was not corrected for B_1_^+^ inhomogeneity because such correction requires non-standard sequence-specific simulations rather than a simple B_1_^+2^ normalization.

The ihMTRAGE files were first concatenated into a 4D volume, then denoised with MP-PCA using MRtrix3’s dwidenoise (Cordero-Grande et al., 2019; Tournier et al., 2019; Veraart, Fieremans, et al., 2016; Veraart, Novikov, et al., 2016), and then iteratively motion-corrected (Soustelle et al., 2020). The iterative motion correction procedure involved looping over the five volumes in the ihMTRAGE acquisition, applying N4 bias correction to each, skull stripping the bias-corrected image with SynthStrip (Hoopes et al., 2022), rigid-body aligning each skull-stripped image to the reference image, and then updating the reference image by averaging it with the newly-registered image. The skull-stripped reference image was used to calculate a transform to T_1_w space.

ihMTw was calculated as the difference between the sum of single-offset (positive and negative) saturated volumes and the sum of dual-offset saturated volumes; ihMTR was then obtained by dividing this ihMTw signal by the unsaturated reference (M0) volume (Alsop et al., 2023; Prevost et al., 2016). MTR was calculated from the single-positive-offset saturated volume as one minus its ratio to the unsaturated reference volume (Van Obberghen et al., 2018). The remaining ihMTRAGE derivatives—ihMTsat and ihMTsat-B_1_c—were estimated using a modified version of the ihmt_proc Python package, which fits the MT pulse saturation parameter from the saturated volumes with corrections derived from the MP2RAGE-based T1 map and the TB1TFL-derived B_1_^⁺^ map (Helms et al., 2008; Munsch et al., 2021).

#### 2.4.6. Diffusion MRI Preprocessing

dMRI preprocessing was performed using QSIPrep 26.0.0 (Cieslak et al., 2021), which is based on Nipype 1.9.1 (Esteban et al., 2025; K. Gorgolewski et al., 2011). The following paragraphs contain partially-edited text generated by QSIPrep, which is distributed with a Creative Commons Zero (CC0) license with the expressed intention of being included in manuscripts.

Using anatomical MRI data, a transform from native space to AC-PC orientation was calculated to be subsequently applied to the dMRI data. For participants with multiple T_1_w images, a participant-average T_1_w-reference map was computed by co-registering the two T_1_w images (after INU-correction) using antsRegistration (ANTs 2.4.3). The anatomical reference image was reoriented into AC-PC alignment using a 6-DOF transform extracted from a full affine registration to the MNI template. A full nonlinear registration to the template from AC-PC space was estimated via symmetric nonlinear registration (SyN) using antsRegistration. Brain extraction was performed on the T_1_w image using SynthStrip (Hoopes et al., 2022) and automated segmentation was performed using SynthSeg (Billot et al., 2023) from FreeSurfer version 7.3.1.

Magnitude and phase dMRI data were combined into a complex-valued file, then denoised using the Marchenko-Pastur PCA method implemented in dwidenoise (Cordero-Grande et al., 2019; Tournier et al., 2019; Veraart, Fieremans, et al., 2016; Veraart, Novikov, et al., 2016) with an automatically-determined window size of 3 voxels. After denoising, the complex-valued data were split back into magnitude and phase, and the denoised magnitude data were retained. The mean intensity of each dMRI series was adjusted so that the mean *b* = 0 signal matched across the AP and PA acquisitions. Each pair of images from a session were merged into a single file, as required for subsequent FSL workflows.

FSL’s eddy was used for head motion correction and eddy current correction (Andersson & Sotiropoulos, 2016). Eddy was configured with a *q*-space smoothing factor of 10, a total of 5 iterations, and 1000 voxels used to estimate hyperparameters. A quadratic first level model and a linear second level model were used to characterize eddy current-related spatial distortion. *q*-space coordinates were forcefully assigned to shells. Field offset was attempted to be separated from participant movement. Shells were aligned post-eddy. Eddy’s outlier replacement was run (Andersson et al., 2016). Data were grouped by slice, only including values from slices determined to contain at least 250 intracerebral voxels. Groups deviating by more than 4 standard deviations from the prediction had their data replaced with imputed values.

Distortion correction was performed in two stages. In the first stage, FSL’s TOPUP (Andersson et al., 2003) was used to estimate a susceptibility-induced off-resonance field based on *b*=0 images extracted from the reverse phase-encoded image pairs. The TOPUP-estimated fieldmap was incorporated into the Eddy current and head motion correction interpolation (Andersson et al., 2017). Interpolation after head motion and initial susceptibility distortion correction was performed using the Jacobian modulation method. DRBUDDI (Irfanoglu et al., 2015), part of the TORTOISE (Irfanoglu et al., 2017) software package, was used to perform the second stage of susceptibility distortion correction. A *b* = 0 image and fractional anisotropy image from both phase encoding directions were used together in a multi-modal registration to estimate the susceptibility-induced off-resonance field. Signal intensity was adjusted in the final interpolated images using a method similar to least squares reconstruction (Andersson et al., 2003). The corrected dMRI time-series were aligned to AC-PC orientation, maintaining the 1.7 mm isotropic voxel resolution.

#### 2.4.7. Diffusion MRI Postprocessing

QSIRecon version 26.0.0 was used to execute multiple dMRI reconstruction workflows in parallel, as described below.

##### 2.4.7.1. NODDI Modeling with AMICO

A neurite orientation dispersion and density imaging (NODDI) model (H. Zhang et al., 2012) was fitted using the AMICO (version 2.0.3) implementation (Daducci et al., 2015). Separate NODDI fits were performed using tissue-specific fixed intrinsic parallel diffusivities for white and gray matter. For the white matter fit, the intrinsic parallel diffusivity was set to 1.7×10^-3^ mm^2^/s, whereas for the gray matter fit it was reduced to 1.1×10^-3^ mm^2^/s, consistent with prior work optimizing NODDI for cortical gray matter (Guerrero et al., 2019). The isotropic diffusivity was fixed at 3.0×10^-3^ mm^2^/s for both fits.

NODDI produced estimates of intracellular volume fraction (ICVF), orientation dispersion index (ODI; output as OD), and isotropic volume fraction (ISOVF). Under the assumptions of the NODDI model, ICVF indexes the relative intra-neurite signal fraction and is commonly interpreted as a proxy for neurite density, ODI characterizes the dispersion of neurite orientations, and ISOVF indexes the isotropic or free-water signal fraction. The pipeline also produced tissue fraction modulated (weighted by 1 - ISOVF) ICVF and ODI maps (Parker et al., 2021). ICVF and ISOVF from the white matter NODDI model were additionally used in the calculation of *g*-ratio maps.

##### 2.4.7.2. Diffusion Kurtosis Modeling with DIPY

DIPY (version 1.10.0) was used to fit a diffusion kurtosis imaging (DKI) model (Henriques et al., 2021), which extends the diffusion tensor model to characterize non-Gaussian diffusion. The fit produced conventional diffusion-tensor measures, including fractional anisotropy (FA), mean diffusivity (MD), axial diffusivity (AD), and radial diffusivity (RD), together with kurtosis-derived measures including mean kurtosis (MK), mean kurtosis tensor (MKT), axial kurtosis (AK), radial kurtosis (RK), and kurtosis fractional anisotropy (KFA). Additional descriptors of diffusion-tensor shape, including linearity, planarity, and sphericity, were also generated. QSIRecon’s DKI implementation produces these tensor and kurtosis outputs from the same DKI fit.

White Matter Tract Integrity (WMTI) (Fieremans et al., 2013) measures were additionally estimated from the DKI model. WMTI interprets the diffusion signal using restricted and hindered diffusion compartments that, under the assumptions of the model and particularly in relatively coherently oriented white matter, are associated with intra-axonal and extra-axonal water, respectively. The compartment-specific outputs included axonal water fraction (AWF), which estimates the fraction of water assigned to the restricted compartment; axonal diffusivity (AxonalD), defined in the DIPY implementation as the trace of the restricted diffusion tensor; and the axial and radial diffusivities of the hindered extra-axonal compartment (ADE and RDE). Extra-axonal tortuosity was calculated as the ratio of ADE-to-RDE.

##### 2.4.7.3. MAP-MRI and Diffusion Tensor Modeling with TORTOISE

TORTOISE (Irfanoglu et al., 2017) software (version 26.1.3) was used to estimate both diffusion tensor and Mean Apparent Propagator MRI (MAP-MRI) models (Özarslan et al., 2013). A diffusion tensor was estimated from both the lower-*b* inner shells of data (*b* < 1200) and the full set of shells, using weighted linear least-squares (WLLS) estimation. These tensor fits produced fractional anisotropy (FA), mean diffusivity (MD), axial diffusivity (AD), radial diffusivity (RD), and lattice index (LI) measures.

MAP-MRI was used to estimate the three-dimensional ensemble average propagator (EAP), providing measures that characterize the displacement distribution without assuming Gaussian diffusion. Derived maps included overall non-Gaussianity (NG), together with non-Gaussianity parallel (NG||) and perpendicular (NG⊥) to the principal diffusion direction; propagator anisotropy (PA); and the angular dissimilarity underlying propagator anisotropy (PAθ) between the estimated propagator and its isotropic counterpart. MAP-MRI additionally produced return-to-origin probability (RTOP), return-to-axis probability (RTAP), and return-to-plane probability (RTPP), which respectively characterize the EAP at zero displacement in three, two, and one dimensions and are sensitive to different aspects of diffusion restriction.

##### 2.4.7.4. Generalized q-Sampling Imaging with DSI Studio

DSI Studio (version “Hou” - commit 94b9c79, Yeh, 2025) was used to perform both diffusion tensor and generalized *q*-sampling imaging (GQI) reconstruction (Yeh et al., 2010). An ordinary least-squares diffusion tensor was fitted using the lower-*b* inner shells, producing fractional anisotropy (FA), mean diffusivity (MD), axial diffusivity (AD), and radial diffusivity (RD) maps.

GQI was performed with a diffusion sampling length ratio of 1.25. GQI reconstructs the spin distribution function (SDF), an ODF-like representation of the distribution of diffusion orientations. Derived scalar measures included generalized fractional anisotropy (GFA), which quantifies angular anisotropy of the SDF; quantitative anisotropy (QA), which characterizes the magnitude of anisotropic SDF peaks above the isotropic background; and ISO, which characterizes the isotropic component of the SDF.

##### 2.4.7.5. Bundle Reconstruction and Tractometry

Fiber response functions were estimated using the “dhollander” algorithm (Dhollander et al., 2016, 2019) in MRtrix3. These response functions were used to estimate fiber orientation distributions (FODs) using multi-shell multi-tissue constrained spherical deconvolution (Jeurissen et al., 2014). The resulting FODs were converted from the MRtrix3 spherical harmonic convention to the convention used by DSI Studio and provided as input to DSI Studio’s AutoTrack function. AutoTrack was used to reconstruct 96 participant-specific white matter bundles across the brain, of which 91 were reconstructed in all included sessions and retained for downstream analyses (**Supplementary Table S2**). Bundle-level macrostructural shape measures (Yeh, 2020) were calculated for each tract. In addition, scalar microstructural maps were summarized within each bundle, including participant-, session-, and metric-specific mean and median values.

#### 2.4.8. Multi-Echo Spin Echo Processing

The MESE scans were used to estimate R_2_ maps. The four MESE echoes were acquired through separate excitations, so each echo was skull-stripped with SynthStrip and registered to the skull-stripped first echo image with ANTs. The root mean squared (RMS) of the four echoes was calculated from the motion-corrected volumes, skull-stripped with SynthStrip, and nonlinearly coregistered to T_1_w space image. R_2_ maps were estimated by fitting a monoexponential decay curve to the MESE data using nonlinear model fitting. The resulting R_2_ maps were then transformed to T1w and MNI spaces. While we acquired one MESE image with a reversed phase encoding direction (PA) for the sake of susceptibility distortion correction, we chose not to use them because some participants lacked the PA acquisition, and instead used a SyN-based nonlinear registration to the T_1_w space image to account for the distortion.

#### 2.4.9. Quantitative Susceptibility Mapping

The MEGRE scans were used to estimate quantitative susceptibility maps using two pipelines: SEPIA and the χ-separation toolbox. Because the first MEGRE echo had a different receiver bandwidth from the remaining echoes, each QSM pipeline was run twice: once using all five echoes (“E5”) and once using only echoes 2–5 (“E4”). The root mean squared of the five echoes was calculated as a reference image, skull-stripped with SynthStrip, and then coregistered to T_1_w space.

E4 and E5 R_2_* maps were estimated by fitting a monoexponential decay curve to the MEGRE data using nonlinear model fitting. E4 and E5 R_2_’ maps were then calculated as the MEGRE-derived R_2_* maps minus the MESE-derived R_2_ map. These R_2_* and R_2_’ maps were transformed to T_1_w and MNI spaces, and were retained as myelin-sensitive relaxation derivatives.

##### 2.4.9.1. Standard QSM Pipeline with SEPIA

We used the SEPIA toolbox to estimate quantitative susceptibility maps from the MEGRE data. Our configuration used BET (Smith, 2002) for brain mask extraction, ROMEO (Dymerska et al., 2021) for phase unwrapping, V-SHARP (Li et al., 2011) for background field removal, and dipole inversion for QSM (Karsa et al., 2019; Schweser et al., 2013). SEPIA-generated QSM maps were computed with CSF as reference. This produced two outputs: “QSM-SEPIA-E5-χ” and “QSM-SEPIA-E4-χ”.

##### 2.4.9.2. χ-Separation QSM Pipeline

We used the χ-separation toolbox (Kim et al., 2025; Shin et al., 2021) to estimate separate paramagnetic and diamagnetic susceptibility maps. The χ-separation toolbox fits a two-pool model using a deep neural network, generating separate paramagnetic and diamagnetic maps, along with a magnetic susceptibility map. The paramagnetic, diamagnetic, and χ maps generated by the toolbox were normalized to CSF.

As we acquired low-resolution MESE data, from which we were able to estimate R_2_ maps, we ran the χ-separation pipeline three ways. In the first configuration, we used the χ-separation toolbox’s built-in R_2_* estimation pipeline, which uses the ARLO algorithm (Pei et al., 2015), to estimate an R_2_* map, from which an R_2_’ map was estimated by applying a scaling factor. We refer to this as the “R_2_*” variant. In the second configuration, the ARLO-estimated R_2_* map was used to estimate an R_2_’ map using the R_2_p-net neural network. We refer to this as the “R_2_pnet” variant. Finally, we provided the MESE and MEGRE-derived R_2_’ map as an input to the χ-separation toolbox. We refer to this as the “R_2_’” variant. Combining the three R_2_ variants with the E4 and E5 echo sets yielded six χ-separation configurations. Each configuration produced paramagnetic, diamagnetic and total susceptibility maps, for a total of 18 susceptibility maps (**Table 2**).

**Table 2.** QSM processing configurations and resulting susceptibility components.

| Name | Software | Number of echoes | Variant | Chi ($\chi$) | Diamagnetic (Dia) | Paramagnetic (Para) |
| --- | --- | --- | --- | --- | --- | --- |
| QSM-SEPIA-E5-χ | SEPIA | 5 | | ✓ | | |
| QSM-SEPIA-E4-χ | SEPIA | 4 | | ✓ | | |
| QSM-χ-R<sub>2</sub><sup>*</sup>-E5 | χ-separation | 5 | R<sub>2</sub><sup>*</sup> | ✓ | ✓ | ✓ |
| QSM-χ-R<sub>2</sub><sup>*</sup>-E4 | χ-separation | 4 | R<sub>2</sub><sup>*</sup> | ✓ | ✓ | ✓ |
| QSM-χ-R<sub>2</sub>pnet-E5 | χ-separation | 5 | R<sub>2</sub>pnet | ✓ | ✓ | ✓ |
| QSM-χ-R<sub>2</sub>pnet-E4 | χ-separation | 4 | R<sub>2</sub>pnet | ✓ | ✓ | ✓ |
| QSM-χ-R<sub>2</sub>'-E5 | χ-separation | 5 | R<sub>2</sub>' | ✓ | ✓ | ✓ |
| QSM-χ-R<sub>2</sub>'-E4 | χ-separation | 4 | R<sub>2</sub>' | ✓ | ✓ | ✓ |

#### 2.4.10. *g*-Ratio Processing

The microscopic *g*-ratio is defined as the ratio of the inner axonal diameter to the total outer diameter of a myelinated fiber. MRI provides an aggregate voxelwise estimate of this quantity based on the relative axonal and myelin volume fractions. We calculated two versions of the *g*-ratio (Stikov et al., 2015), using ihMTsat-B_1_c and ihMTR measures from the ihMTRAGE data as two proxies for myelin volume fraction (MVF), and the ISOVF and ICVF measures from the NODDI outputs to calculate the axonal volume fraction (Berg et al., 2022). First, we warped the ISOVF and ICVF images from AC-PC space to T1w space (i.e., from QSIRecon’s output anatomical space to fMRIPrep’s output anatomical space). Next, we derived a splenium mask from the FreeSurfer aseg image and warped it from fsnative space to T_1_w space at the same resolution. We then extracted the mean value within the splenium from the ISOVF, ICVF, ihMTR, and ihMTsat-B_1_c images for each participant and session. These values were collected into a tabular file. Once collected, we computed the scaling factor such that the mean *g*-ratio in the splenium across the full sample would be 0.7. This resulted in a scaling factor of ∼2.214 for the ihMTR-derived MVF and ∼0.097 for the ihMTsat-B_1_c-derived MVF. The algebraic derivation of the scaling factor used for *g*-ratio calibration is provided in **Supplementary Note S1**. The associated *g*-ratio maps were calculated for each of the participants and sessions, in T_1_w space with 1 mm^3^ isotropic voxels.

#### 2.4.11. *q*-Ratio Processing

The *q*-ratio is defined as the ratio of R_1_ to T_2_* (equivalent to R_1_ x R2*), and has been proposed as a myelin sensitive derivative (Haast et al., 2016). The *q*-ratio has previously exhibited better contrast between myelinated and demyelinated regions than either R_1_ or R_2_* on their own (Kawaguchi et al., 2024). We calculated four versions of the *q*-ratio (**Table 3**), using MEGRE-derived R_2_* and MP2RAGE-derived R_1_ measures, based on the procedure in (Kawaguchi et al., 2024). We used each combination of the E4 and E5 R_2_* scalars and the R_1_ and B_1_-corrected R1 scalars. The *q*-ratio maps were calculated in T_1_w space and were then warped to MNI space.

**Table 3.** ***q*-Ratio processing configurations.**

| Name | Number of MEGRE echoes | MP2RAGE variant |
| --- | --- | --- |
| *q*-Ratio-E5 | 5 | R<sub>1</sub> |
| *q*-Ratio-E5-B<sub>1c</sub> | 5 | R<sub>1</sub>-B<sub>1c</sub> |
| *q*-Ratio-E4 | 4 | R<sub>1</sub> |
| *q*-Ratio-E4-B<sub>1c</sub> | 4 | R<sub>1</sub>-B<sub>1c</sub> |

### 2.5. Myelin-Sensitive Map Analyses

For the primary analyses, we focused on a subset of 27 myelin-sensitive metrics selected to span the major processing workflows, avoid closely related redundant variants, and reflect their use in prior myelin imaging studies (**Figure 1**). All analyses were also repeated using a broader panel of 93 derived metrics (**Supplementary Table S3**), with these results presented throughout the **Supplementary Materials**. We first quantified the degree to which each metric differentiated gray and white matter. We then assessed relationships among metrics using intermetric correlations and evaluated test–retest reliability using intraclass correlation coefficients (ICCs). Finally, we calculated discriminability to quantify the extent to which spatial profiles from the same participant were more similar across sessions than profiles from different participants (Bridgeford et al., 2021). Metrics were evaluated separately across tissue compartments (white and gray matter) and spatial scales (voxelwise and regionwise) where applicable. For ICVF, gray matter analyses used the gray matter-optimized NODDI fit, whereas white matter analyses used the conventional white matter NODDI fit. The *g*-ratio was analyzed only within white matter, where the assumption that NODDI-derived ICVF primarily reflects axonal populations is more appropriate than in gray matter.

**Figure 1.**
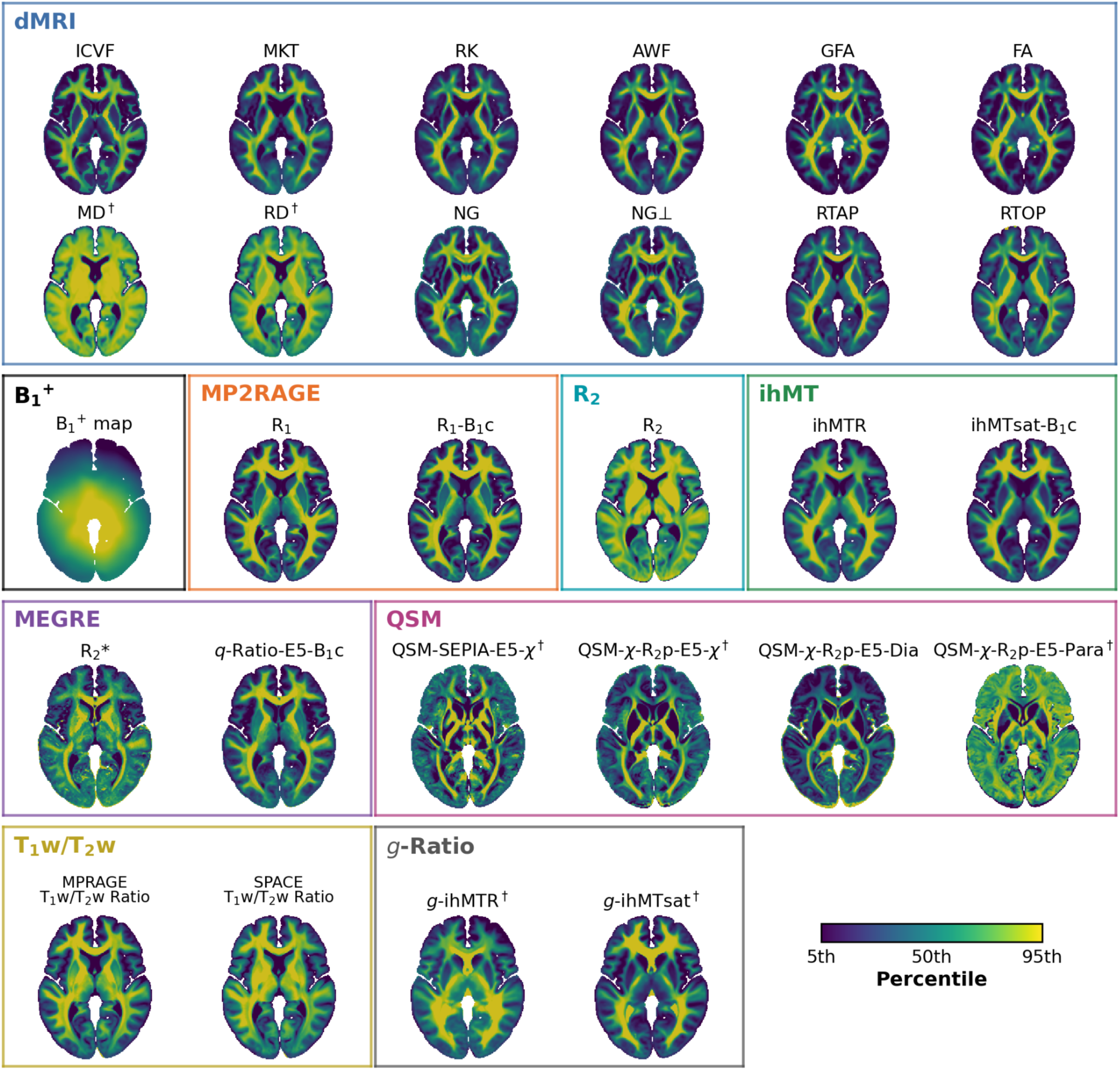
Myelin-sensitive maps used in primary analyses. The same axial slice in the middle of the brain is shown for all maps. Each map depicted is the average MNI-space image across all sessions that passed quality control. Metrics are grouped by their metric family. Each metric is displayed using its own intensity scale spanning the 5th to 95th percentiles of the corresponding group-average map, with the median indicated at the midpoint. The *g*-ratio was only analyzed within white matter. The ICVF image shown here is spliced such that the white matter and gray matter values are derived from the two corresponding NODDI models. The average B_1_^+^ bias map, which is used to correct for bias inhomogeneities, is also depicted. Metrics marked by dagger symbols (†) indicate the colormap was inverted such that values would appear brighter in white matter for consistency across metrics.

#### 2.5.1. Quality Assurance

We generated participant-and session-level quality-assurance reports to visually inspect metric-wise quality and spatial co-registration. Visual inspection identified no substantial misregistration between the metric maps and their corresponding anatomical references. We identified 16 source images with visually apparent artifacts whose derivatives were excluded from subsequent analyses **(Figure 2)**.

**Figure 2.**
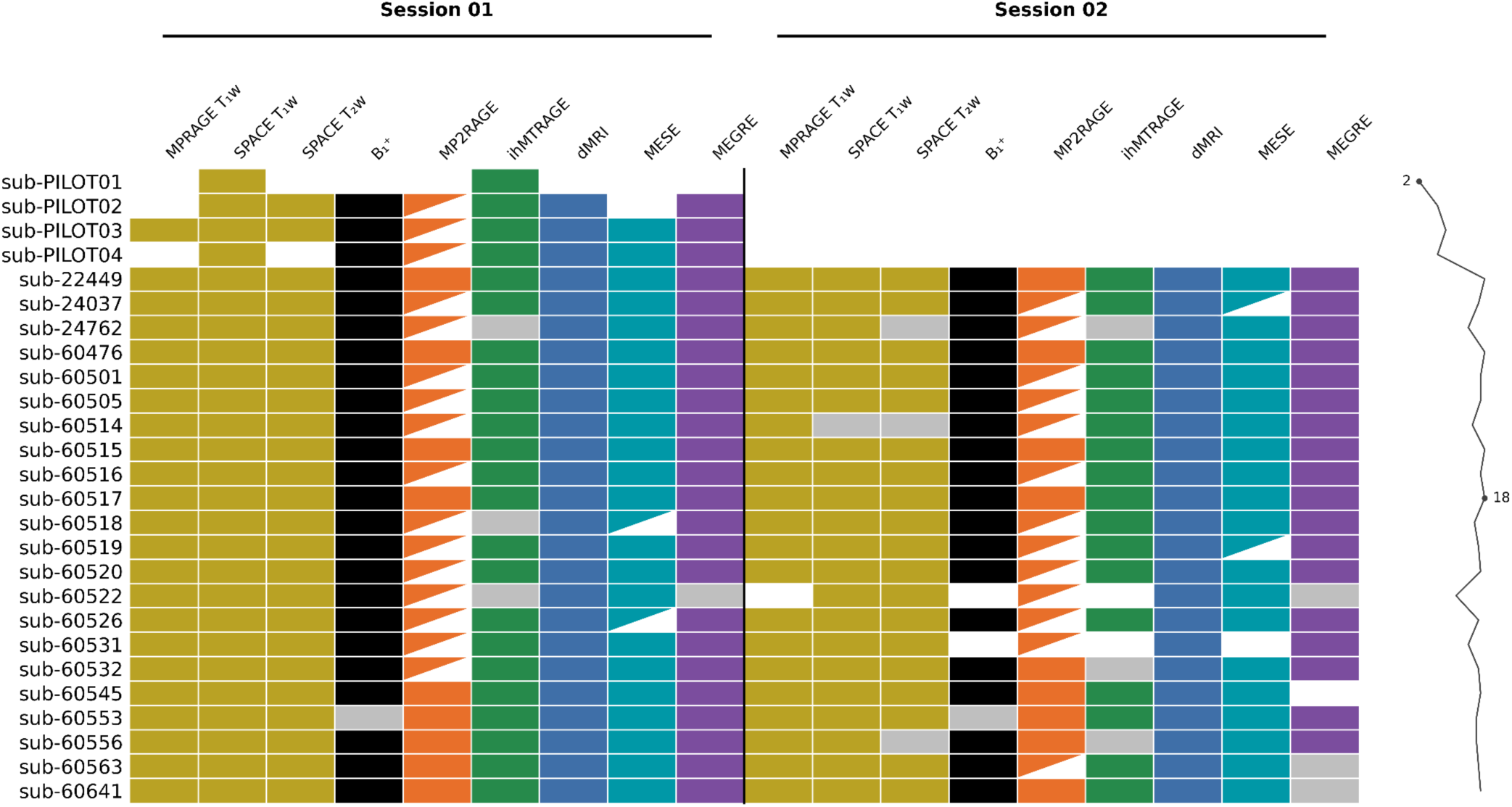
Study Completion by Participant. The second session was terminated early for three participants, resulting in missing acquisitions. Partial cells in the MP2RAGE columns refer to scans without reconstructed phase data. Partial cells in the MESE columns refer to scans without a corresponding reverse phase-encoded acquisition. Cells that are gray indicate scans that were acquired, but which failed manual quality control. The black line to the right reflects how many acquisitions are present and pass quality assurance for each participant.

#### 2.5.2. Tissue Compartment Definition and Parcellation

For white matter region-wise analyses, metrics were summarized within the 91 participant-specific bundles that were consistently reconstructed across all analyzed participants and sessions by the QSIRecon tractography workflow. For non-dMRI metrics, reconstructed bundles were registered from AC-PC to T1w space. For each participant, session, metric, and bundle, the median metric value was retained for subsequent analyses.

Gray matter region-wise metrics were summarized using participant-specific Desikan– Killiany–Tourville (DKT) cortical parcellations (Desikan et al., 2006; Klein & Tourville, 2012) generated by FreeSurfer on the participant-averaged T_1_w image. The DKT atlas comprises 31 cortical regions per hemisphere. DKT labels were additionally registered to AC-PC space using the T_1_w-to-ACPC transforms generated by QSIPrep with “genericLabel” interpolation. For each participant, session, metric, and cortical parcel, the median metric value was retained for subsequent analyses.

Voxelwise analyses were performed in MNI space. To provide a voxelwise gray matter compartment comparable to the cortical regions represented by the DKT atlas, gray matter analyses were restricted to the cortical ribbon. We used two tissue-masking strategies depending on whether a common mask across participants was required for the particular test, as described below. For all analyses, MNI-space metric maps were re-gridded to the tissue segmentation image with linear interpolation.

For voxelwise ICC analyses, which estimate reliability across participants at each voxel, fixed group-level gray and white matter masks were required so that ICCs at a given voxel were estimated from the same anatomical location across participants. These were defined using the 1mm isotropic resolution “aseg_dseg” segmentation in MNI space from Templateflow (Ciric et al., 2022). For all analyses besides ICC calculation, participant-specific tissue masks were used to afford improved anatomical specificity. For these remaining analyses, statistics were first calculated within each participant and subsequently pooled across participants. White matter was defined using each participant’s deterministic SynthSeg white matter segmentation. We transformed the gray matter ribbon output from fMRIPrep’s anatomical preprocessing workflow to MNI space with “genericLabel” interpolation.

#### 2.5.3. Gray–White Matter Differentiation

To quantify how strongly each metric differentiated gray and white matter, we compared voxelwise metric distributions between the two tissue compartments. Gray and white matter were defined using participant-specific tissue segmentations in MNI space. Within each tissue, finite, nonzero values were retained after excluding extreme values using a robust median absolute deviation (MAD) *z* score threshold of 6. For each participant and session, gray–white matter separation was quantified using a robust standardized median difference, calculated as the difference between the white matter and cortical gray matter medians divided by a pooled MAD-based estimate of dispersion:

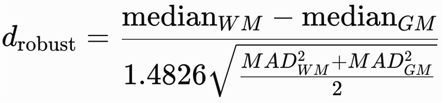

The factor 1.4826 scales the MAD to be comparable to the standard deviation under a normal distribution. Session-level effect sizes were averaged within participants, and the group-level effect size for each metric was calculated as the mean across participants with available data. Positive values indicate higher metric values in white matter than cortical gray matter. Uncertainty in the group mean was quantified using 10,000 participant-level bootstrap resamples with replacement; 95% confidence intervals were defined by the 2.5th and 97.5th percentiles of the bootstrap distribution.

#### 2.5.4. Intermetric Correlations

To evaluate similarity among myelin-sensitive metrics, we calculated intermetric spatial correlations. For voxelwise analyses, voxels were considered eligible for a given metric only when values were finite and nonzero. To reduce the influence of extreme values, metric-specific outliers were additionally excluded using a robust *z* score based on the tissue-specific median and median absolute deviation (MAD), with values exceeding an absolute robust *z* score of 6 removed. For each pair of metrics, Pearson’s correlation coefficient was calculated across the intersection of their valid voxel masks within each tissue compartment. Correlation coefficients were Fisher *z*-transformed and averaged across available sessions within each participant and then across participants, giving each participant equal weight in the group-level estimate regardless of the number of contributing sessions. The resulting group-average *z* values were transformed back to correlation coefficients. Hierarchical clustering of the voxelwise correlation matrices was performed separately within white and gray matter using absolute-correlation distance (1-|*r*|) and average linkage.

To assess whether intermetric relationships were preserved across tissue compartments, we additionally quantified the similarity of each metric’s correlation profile between white and cortical gray matter. For each metric present in both tissue-specific matrices, we extracted its correlations with all other shared metrics, excluding the self-correlation, separately in white and gray matter. Pearson’s correlation coefficient was then calculated between the white-and gray-matter correlation profiles, with larger values indicating greater preservation of that metric’s pattern of intermetric relationships across tissue compartments.

For supplementary region-wise analyses, Pearson’s correlation coefficient was calculated across metric profiles within gray matter parcels or white matter bundles using regions with finite values for both metrics. Regional correlation coefficients were averaged across sessions and participants using the same Fisher z procedure described above.

#### 2.5.5. Test–Retest Reliability

Test–retest reliability was assessed using the intraclass correlation coefficient, ICC(2,1), corresponding to a two-way random-effects model of absolute agreement for a single measurement. ICC values were calculated independently for each metric at each voxel, gray matter parcel, and white matter bundle. Participants were included for a particular metric’s ICC measurement only when the corresponding source image data from both sessions were available and passed quality assurance. For each metric and tissue compartment, voxelwise or regionwise ICC values were summarized by their median and interquartile range. To aid interpretation of ICC values, we additionally compared within-participant test–retest variability to between-participant variability. Within-participant variability was estimated from the residual mean square of a participant-by-session ANOVA, and was expressed relative to the between-participant standard deviation.

#### 2.5.6. Discriminability

To assess whether each imaging metric captured participant-specific spatial patterns, we calculated discriminability (Bridgeford et al., 2021) for gray matter parcels and white matter bundles. Higher discriminability indicates that a metric captures spatial patterns that are reproducibly more similar within an individual than between individuals, making it potentially useful for studies of individual differences or participant-specific alterations. Lower discriminability indicates less participant-specific spatial information, which may reflect spatial patterns that are largely shared across individuals or greater measurement variability. For each metric, participant-session profiles were constructed from parcel-or bundle-wise values, and each regional feature was *z*-scored across profiles. Pairwise distances between profiles were calculated using correlation distance. For each participant-session profile, the distance to the matching profile from the same participant was compared with distances to profiles from all other participants. Discriminability was defined as the proportion of between-participant distances that exceeded the within-participant distance, averaged across profiles, with values approaching 1 indicating greater participant specificity. Discriminability was also calculated using voxelwise profiles; however, all metrics reached the upper bound of discriminability because of the high dimensionality of these profiles.

## 3. Results

### 3.1. Gray–White Matter Differentiation Varies Markedly Across Metrics

Gray–white matter differentiation varied markedly across metrics (**Figure 3**). Most metrics showed higher values in white matter voxels than in gray matter voxels, with the strongest separation observed for the MP2RAGE-derived R_1_ and R_1_-B_1_c measures, followed by ICVF and the MPRAGE T_1_w/T_2_w ratio. Notably, the strong gray–white matter differentiation observed for ICVF may partly reflect the use of separate tissue-specific NODDI fits in gray and white matter. Other measures including ihMTsat-B_1_c, ihMTR, and *q*-Ratio-E5-B_1_c also showed strong white matter–gray matter differentiation, as did several diffusion-derived measures including AWF, MKT, RK, FA, GFA, RTOP, RTAP, NG, and NG⊥. In contrast, RD and MD showed large effects in the opposite direction, reflecting higher values in cortical gray matter than white matter. R_2_ and R_2_* showed more moderate white matter–gray matter separation. QSM-derived measures generally exhibited the weakest tissue differentiation and varied in direction: QSM-χ-R_2_p-E5-Dia was higher in white matter, QSM-χ-R_2_p-E5-χ and QSM-SEPIA-E5-χ were higher in cortical gray matter, and QSM-χ-R_2_p-E5-Para showed little systematic difference between tissue compartments. Overall, the magnitude and direction of gray–white matter differentiation depended strongly on the imaging contrast, with the clearest separation observed for relaxometry-, T1w/T_2_w-, ihMT-, and several diffusion-derived metrics and substantially weaker differentiation among QSM measures. Differentiation profiles for all metrics are reported in **Supplementary Figure S2.**

**Figure 3.**
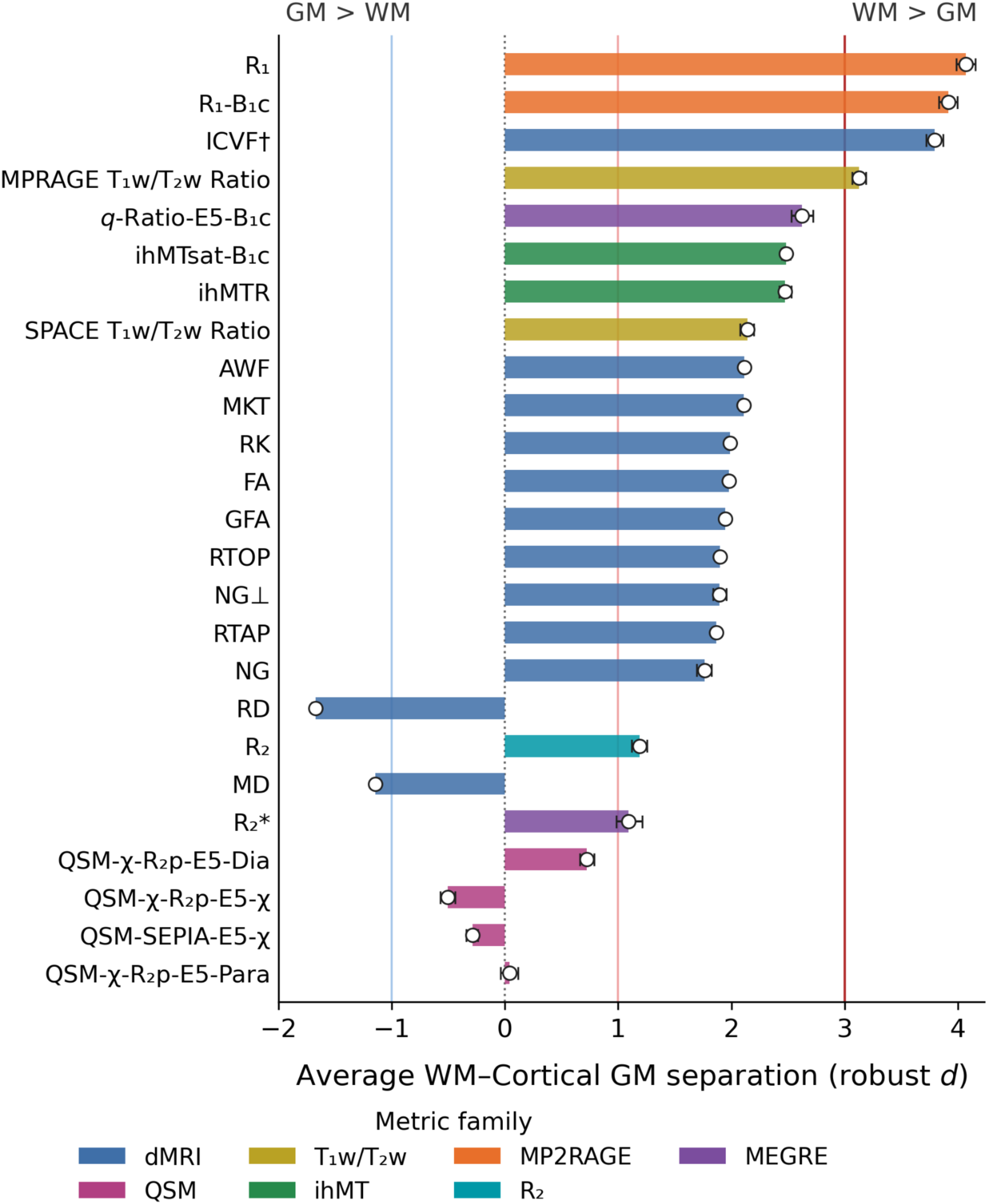
Differentiation of cortical gray and white matter across imaging metrics. Gray–white matter differentiation was quantified voxelwise within each participant and session using a robust standardized median difference, calculated as the difference between the white matter and cortical gray matter medians divided by a pooled MAD-based estimate of dispersion. Session-level effect sizes were averaged within participants and then averaged across participants. Positive values indicate higher metric values in white matter than cortical gray matter. Bars show the mean participant-level effect size for each metric, with error bars representing 95% confidence intervals obtained from 10,000 participant-level bootstrap resamples. Metrics are ordered by the absolute magnitude of their mean effect size. Colors denote metric families. †: For ICVF, white matter and gray matter values were derived from the two tissue-specific NODDI models.

### 3.2. Intermetric Correlations are Strongest within Modeling Families

Intermetric correlations revealed structured spatial relationships that were generally stronger in white matter and among metrics derived from the same acquisition or modeling family (**Figure 4**). In both tissue compartments, QSM-derived metrics tended to show comparatively distinct correlation patterns relative to the other metric families. In white matter voxels (**Figure 4A**), MP2RAGE-and ihMT-derived measures were strongly associated with several dMRI-derived measures, particularly measures derived from diffusion kurtosis and related higher-order diffusion models. In cortical gray matter voxels (**Figure 4B**), RD and MD exhibited especially strong negative spatial relationships with several ihMT-derived measures, whereas RTOP and RTAP showed corresponding positive relationships.

**Figure 4.**
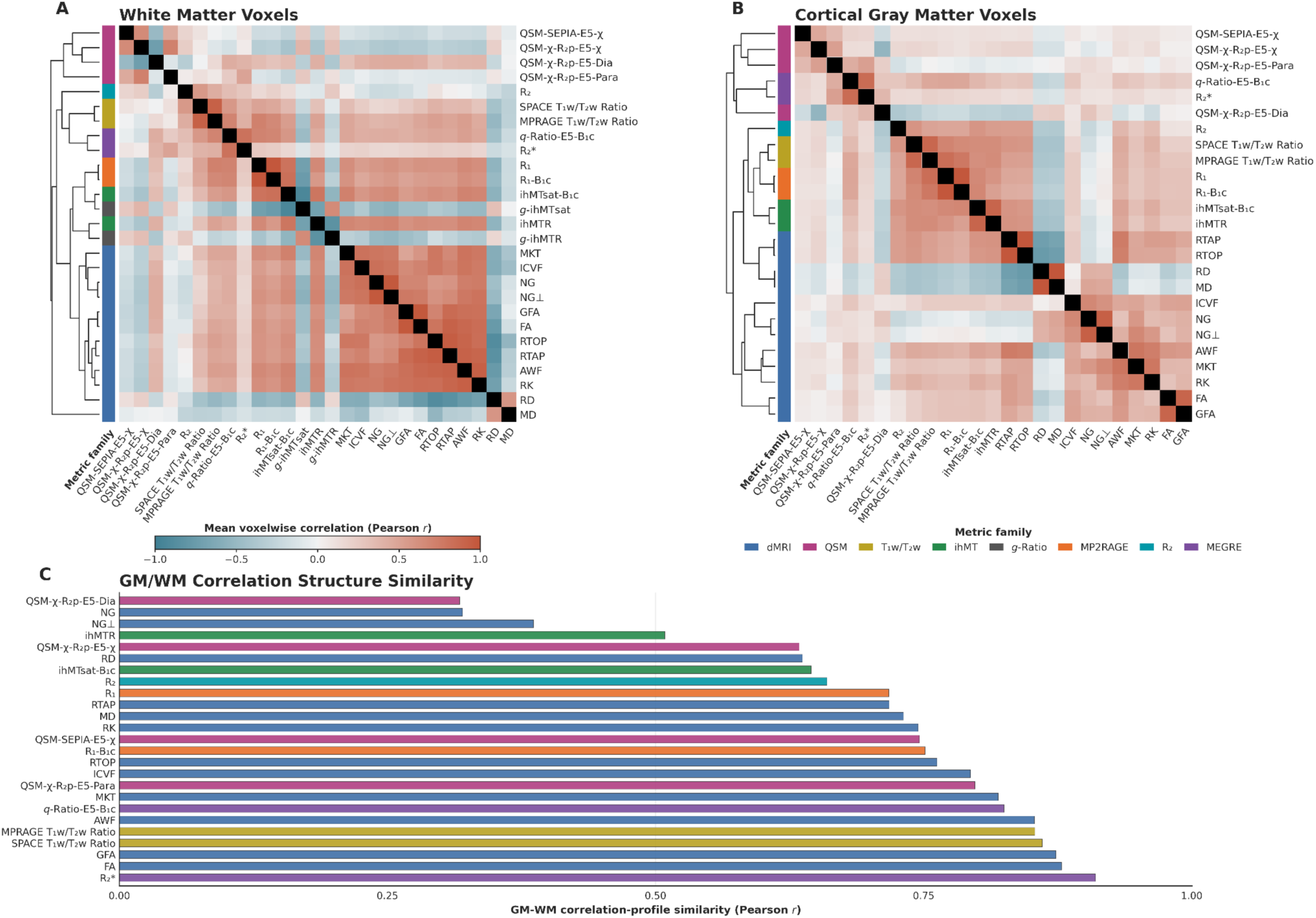
Intermetric spatial correlations across white and gray matter. Intermetric correlation matrices are shown for **(A)** white matter voxels and **(B)** cortical gray matter voxels. Matrices show Pearson correlations calculated across pairwise-valid voxels within each participant and session. Correlation coefficients were Fisher z-transformed and averaged across sessions within participants and then across participants before transformation back to correlation coefficients. Colored bars denote metric families. Hierarchical clustering was performed separately for the white-and gray-matter matrices using absolute-correlation distance and average linkage. **(C)** Similarity of intermetric correlation structure between gray and white matter. For each metric shared between tissue compartments, its correlations with all other shared metrics were extracted from the corresponding white-and gray-matter matrices, excluding the self-correlation. Pearson correlation between these two profiles was used to quantify gray–white matter correlation-profile similarity, with larger values indicating greater preservation of intermetric relationships across tissue compartments. Bars are colored according to metric family.

The extent to which these intermetric relationships were preserved between gray and white matter varied considerably across metrics (**Figure 4C**). Several measures, including R_2_*, FA, GFA, T_1_w/T2w ratios, and multiple advanced dMRI-derived metrics, showed relatively similar correlation profiles across tissue compartments, indicating that their relationships with the broader metric set were largely preserved between gray and white matter. In contrast, several QSM-derived measures, dMRI non-Gaussianity, and ihMTR showed lower gray–white matter profile similarity, suggesting greater tissue dependence in their intermetric relationships. Voxel-, parcel-, and bundle-wise correlation profiles for all metrics are reported in **Supplementary Figures S3-S6**.

### 3.3. Test–Retest Reliability is Generally Strong among Metrics

Test–retest reliability varied substantially across metrics, although most measures showed high reliability across tissue compartments and spatial scales **(Figure 5)**. dMRI-derived measures showed consistently high reliability, with many median ICCs approaching or exceeding 0.90 across tissue compartments and spatial scales. T_1_w/T_2_w and MP2RAGE-derived R_1_ measures also generally exhibited high reliability across tissue compartments. In contrast, QSM-, ihMT-, and R_2_*-derived measures showed greater variability in reliability. R_2_* exhibited particularly low reliability in white matter voxels, while ihMT-derived measures showed reduced reliability across gray matter parcels. The comparatively low cortical gray matter ICCs of the ihMT-derived measures were in part driven by high within-participant test–retest variability compared to between-participant variability (**Supplementary Figure S7**). Expanded ICC summaries for voxels, bundles, and parcels are reported in **Supplementary Figures S8–S11**.

**Figure 5.**
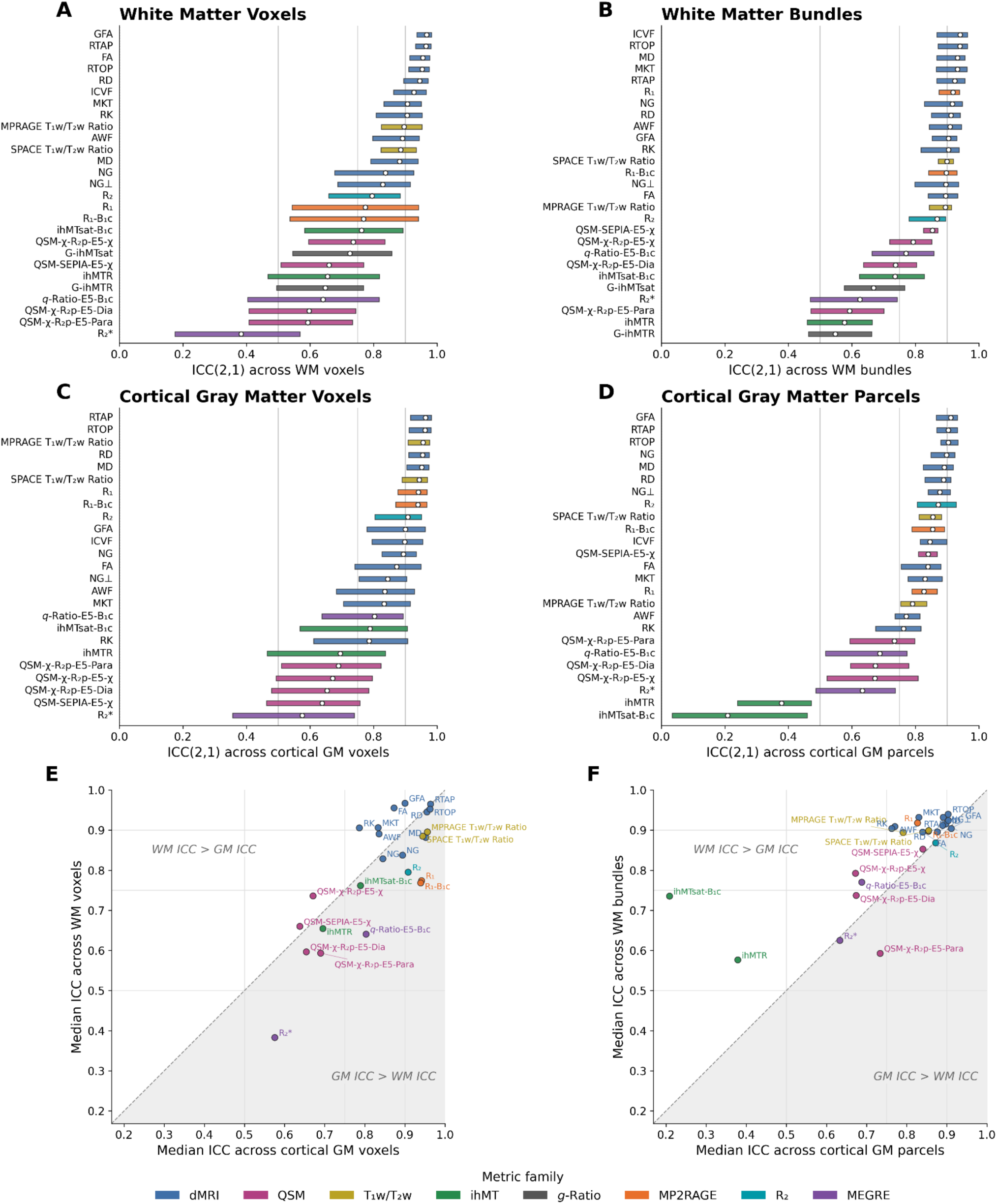
Test–retest reliability of myelin-sensitive metrics across tissue compartments and spatial scales. Test–retest reliability was quantified using ICC(2,1) and summarized across **(A)** white matter voxels; **(B)** white matter bundles; **(C)** gray matter voxels; **(D)** gray matter parcels. White circles indicate the median ICC and the colored bar spans the interquartile range across voxels, bundles, or parcels. **(E–F)** Median ICC values for matched metrics in gray and white matter for voxelwise and regional analyses. Colors denote metric families.

### 3.4. Regional Spatial Profiles are Highly Participant-Specific

Discriminability was uniformly high across metrics, with perfect discriminability (1.00) observed for 18 of 27 metrics in white matter bundles and 14 of 25 metrics in gray matter parcels (**Table 4**). Discriminability reached its upper bound (1.00) for all metrics when using voxelwise profiles, consistent with the high dimensionality of these data. Thus, participant-specific regional spatial profiles were highly distinguishable for nearly all metrics, with appreciable reductions in discriminability largely restricted to R_2_* and ihMT-derived measures, particularly in gray matter. Discriminability values across the expanded supplementary metric set are reported in **Supplementary Figure S12**.

**Table 4.** Discriminability of white matter bundle and cortical gray matter parcel profiles. Metric families are ordered by their average ICC across tissue compartments, with lower-reliability families appearing lower in the table. Metrics with identical white matter and gray matter discriminability values are combined within each metric family. The asterisk (*) indicates that ICVF values in white matter were derived from the standard NODDI model and gray matter values were derived from a separate gray matter NODDI model.

| Metric Name | Discriminability (WM) | Discriminability (GM) |
| --- | --- | --- |
| **dMRI** |  |  |
| FA, GFA, ICVF*, MD, MKT, NG, NG$\perp$, RD, RTAP, RTOP | 1 | 1 |
| AWF | 1 | 0.997 |
| RK | 1 | 0.996 |
| **MP2RAGE** |  |  |
| $R_1$, $R_1$-B$_1$c | 1 | 1 |
| **T$_1$w/T$_2$w** |  |  |
| SPACE T$_1$w/T$_2$w Ratio | 1 | 1 |
| MPRAGE T$_1$w/T$_2$w Ratio | 1 | 0.999 |
| **R$_2$** | 1 | 0.998 |
| **QSM** |  |  |
| QSM-χ-R$_2$p-E5-χ | 1 | 1 |
| QSM-χ-R$_2$p-E5-Para | 0.999 | 0.999 |
| QSM-SEPIA-E5-χ | 0.998 | 0.997 |
| QSM-χ-R$_2$p-E5-Dia | 0.997 | 0.995 |
| **MEGRE** |  |  |
| $q$-Ratio-E5-B$_1$c | 0.983 | 0.928 |
| R$_2^*$ | 0.923 | 0.901 |
| **g-ratio** |  |  |
| $g$-ihMTsat | 0.99 | — |
| $g$-ihMTR | 0.928 | — |
| **ihMT** |  |  |
| ihMTsat-B$_1$c | 0.996 | 0.774 |
| ihMTR | 0.968 | 0.792 |

## 4. Discussion

Non-invasive imaging of myelin has emerged as an important goal for characterizing brain development, aging, plasticity, and disease. However, principled method selection and synthesis across studies remain challenging because many myelin-sensitive MRI techniques have not been systematically compared within the same participants. Here, we introduced and evaluated MIRROR, an openly available multimodal test–retest resource containing a diverse set of MRI acquisitions capable of producing several myelin-sensitive brain maps. Collectively, our findings demonstrate that candidate MRI measures, while broadly reliable and participant-specific, differ meaningfully in their tissue contrast and relationship with one another. Moreover, these intermetric relationships were not fully preserved between gray and white matter, indicating that correspondence among measures is tissue dependent and may further complicate synthesis across studies that focus on different tissue compartments. To facilitate methodological benchmarking and development across the neuroimaging community, we openly release the raw and processed imaging data together with standardized processing code.

Consistent with the different biological substrates influencing each image contrast, gray–white matter differentiation varied considerably across metrics. MP2RAGE-derived R_1_ measures exhibited the strongest differentiation, while QSM-derived measures generally exhibited weaker and more heterogeneous tissue differentiation. This variation is expected because the evaluated contrasts are influenced by distinct tissue properties including macromolecular content, water mobility and content, iron, and microstructural organization (Mancini et al., 2020; Piredda et al., 2021; Rahmanzadeh et al., 2022). Gray and white matter differ substantially across many of these properties, in addition to myelin content. Accordingly, strong gray–white matter separation should not itself be interpreted as evidence of myelin specificity.

Notably, several diffusion measures derived from distinct modeling frameworks showed highly similar spatial organization. For example, MKT, NG, and ICVF broadly clustered together and were positively associated with non-dMRI measures such as ihMT. However, cross-modal correspondence was generally stronger in white matter than in cortical gray matter, potentially reflecting the more coherent axonal organization of white matter compared with the greater heterogeneity of neurite orientations and cellular architecture in the cortex. MKT, NG, and ICVF are derived from advanced dMRI reconstruction approaches that leverage multishell data to characterize non-Gaussian and complex diffusion patterns that cannot be captured by the conventional diffusion tensor model. This convergence parallels recent findings that these advanced dMRI measures were associated with cortical myelin density measured using histology and magnetization transfer imaging in the primate cortex (Reveley et al., 2024). More generally, similarities among these dMRI metrics may reflect the fact that ostensibly distinct diffusion models are sensitive to overlapping components of the underlying diffusion signal; recent work has shown that many advanced diffusion measures can be summarized by a smaller number of latent dimensions capturing distinct aspects of water diffusion, such as overall diffusivity, anisotropy, and non-Gaussian diffusion or tissue complexity (Sadikov et al., 2025). This raises the possibility that combinations of complementary dMRI-derived measures could recover a meaningful component of myelin-related signal without requiring a dedicated myelin-sensitive acquisition. Determining the extent to which such diffusion-derived components reflect myelin, rather than other correlated features of tissue microstructure, will require validation against independent histological references in future work.

Test–retest reliability and discriminability provided complementary perspectives on measurement stability. ICC quantifies the extent to which individual differences at a given voxel or region are preserved across sessions, whereas discriminability evaluates whether the multivariate spatial pattern of a metric remains sufficiently stable to distinguish one participant from others. Most metrics exhibited high discriminability even when their local ICCs were more variable, suggesting that participant-specific information may be distributed across many spatial features rather than depending on highly reliable measurements at every individual location. ihMT measures in gray matter were a notable exception. Their low parcel-wise ICCs reflected relatively high within-participant test–retest variability compared with between-participant variation. ihMT also has particularly low signal-to-noise ratio in gray matter (Taso et al., 2023). ihMTR and ihMTsat-B_1_c correspondingly also exhibited reduced gray matter discriminability. Collectively, these findings suggest that ihMT approaches may be less well suited to cortical gray matter analyses, at least using acquisition times feasible for conventional neuroimaging protocols.

Among the evaluated approaches, dMRI, MP2RAGE, and T_1_w/T_2_w ratio mapping offer several practical advantages for studies seeking scalable approaches to myelin-sensitive tissue characterization. Multishell dMRI supports participant-specific tractography while also enabling the wide range of reconstruction approaches applied here. In MIRROR, diffusion-derived measures were consistently among the most reliable and participant-specific metrics in both gray and white matter. MP2RAGE similarly provides both quantitative T_1_/R_1_ measures and a high-resolution T_1_-weighted anatomical image that can be used in many standard neuroimaging workflows. MP2RAGE-derived R_1_ measures exhibited high reliability and showed the strongest gray–white matter differentiation among the evaluated metrics. T_1_w/T_2w_ ratio mapping has the practical advantage of leveraging anatomical scans that may already be collected for other purposes. While T_1_w images are already ubiquitously acquired in neuroimaging studies, T_2_w-weighted images can improve delineation of the pial surface during cortical surface reconstruction and aid in anatomical registration processes. Thus, these approaches that are often already included in multi-modal imaging protocols may provide favorable combinations of myelin sensitivity and reliability.

Several limitations should be considered when interpreting these findings. First, the sample was modest in size and consisted of healthy young adults; it was not designed to represent the broader population across age, demographic, or clinical dimensions. Second, the present dataset lacks direct histological ground truth. The evaluated MRI measures should therefore be considered myelin-sensitive rather than direct or specific measures of myelin content. Third, several evaluated measures—including R1, R_2_, R_2_*, R_2_′, QSM, *q*-ratio, and ihMT-derived contrasts—may exhibit orientation dependence in white matter, such that observed spatial variation may partly reflect fiber orientation relative to the main magnetic field rather than tissue composition alone (Girard et al., 2017; Kawaguchi et al., 2024; Kor et al., 2019; Morris et al., 2022; Pampel et al., 2015; Tax et al., 2021; Wallstein et al., 2024). Fourth, the source acquisitions differed substantially in native spatial resolution and voxel geometry, with dMRI, ihMT, and MESE acquired at lower spatial resolutions than several of the anatomical and quantitative acquisitions. These differences may influence partial-volume effects and sensitivity to fine anatomical variation. Finally, the selection of MRI sequences reflected practical constraints related to scan time, site-specific implementation, as well as access to specialized acquisition methods and postprocessing tools such as dedicated bias correction algorithms. MIRROR therefore does not exhaustively sample the growing range of myelin-sensitive MRI approaches, including methods that often require longer or more specialized acquisitions such as myelin water imaging and mcDESPOT.

MIRROR provides a foundation for developing and evaluating new myelin-sensitive imaging approaches beyond the individual metrics examined here. Future work can leverage the multimodal design to identify shared latent dimensions across diffusion and other contrasts, test whether these components better capture myelin-related variation, and validate them against histological or complementary MRI-based references. The dataset can also support optimization of more parsimonious acquisition protocols that balance biological sensitivity, reliability, redundancy, and scan time. By providing an openly available multimodal test–retest benchmark, MIRROR can support transparent and reproducible development of more efficient and biologically interpretable approaches to in vivo myelin imaging.

## Data and Code Availability

Code for curating, processing, and analyzing the dataset is available at https://github.com/PennLINC/mirror. The modified versions of ihmt_proc and pymp2rage used for processing are available at https://github.com/PennLINC/ihmt_proc/tree/mirror-dev (forked from https://github.com/lsoustelle/ihmt_proc commit hash 436d92c) and https://github.com/PennLINC/pymp2rage/tree/mirror-dev (forked from https://github.com/Gilles86/pymp2rage commit hash 76178f4), respectively. Derivatives and processing code for each modality are made publicly available on OpenNeuro at the following location: https://openneuro.org/datasets/ds008825.

## Author Contributions

**Steven L. Meisler**: Software, Validation, Formal analysis, Writing - Original Draft, Writing - Review & Editing, Visualization. **Taylor Salo**: Conceptualization, Methodology, Software, Validation, Formal analysis, Data Curation, Writing - Original Draft, Writing - Review & Editing, Visualization. **Matthew Cieslak**: Methodology, Software, Writing - Review & Editing. **Brooke L. Sevchik**: Investigation, Project administration, Writing - Review & Editing. **Juliette Brook**: Investigation, Project administration, Writing - Review & Editing. **Guillaume Duhamel**: Methodology, Resources, Writing - Review & Editing. **Olivier M. Girard**: Methodology, Resources, Writing - Review & Editing. **Bart Larsen**: Writing - Review & Editing. **Shraddha Pandey**: Methodology, Software, Writing - Review & Editing. **Lucinda M. Sisk**: Writing - Review & Editing. **Lucas Soustelle**: Methodology, Resources, Writing - Review & Editing. **Valerie J. Sydnor**: Writing - Review & Editing. **Poljanka Johnson**: Writing - Review & Editing. **David R. Roalf**: Writing - Review & Editing. **Russell T. Shinohara**: Writing - Review & Editing. **Manuel Taso**: Conceptualization, Methodology, Resources, Supervision, Writing - Review & Editing. **M. Dylan Tisdall**: Conceptualization, Methodology, Resources, Supervision, Writing - Review & Editing. **Theodore D. Satterthwaite**: Conceptualization, Resources, Supervision, Project administration, Funding acquisition, Writing - Original Draft, Writing - Review & Editing.

## Ethics

All study protocols were approved by the University of Pennsylvania Institutional Review Board (Protocol 857664 / 24-2318). All participants provided written informed consent to enroll in the study.

## Acknowledgements

The authors thank Adam Raikes and Ariel Rokem for their guidance on derivatives calculations. We thank the Center for Advanced Magnetic Resonance Imaging and Spectroscopy at the University of Pennsylvania (RRID: SCR_022398) for their assistance with the acquisition of MRI data.

