## Supplement for "An Open Test–Retest Dataset of Rapid Myelin Mapping Acquisitions"

### Supplementary Materials for Meisler*, Salo*, et al.

| Scalar(s) | Source Acquisitions | Total Duration (MM:SS) |
|---|---|---|
| dMRI | dMRI<br>MPRAGE $T_1$w | 12:48 + 4:05 (16:53) |
| $R_1$ | MP2RAGE | 7:59 |
| $R_1$-$B_1$c | MP2RAGE<br>$B_1^+$ field map | 7:59 + 0:21 (8:20) |
| ihMTw<br>ihMTR<br>MTR | ihMTRAGE<br>MPRAGE $T_1$w | 8:22 + 4:05 (12:27) |
| ihMTsat<br>ihMTsat-$B_1$c | ihMTRAGE<br>$B_1^+$ field map<br>MP2RAGE | 8:22 + 0:21 + 7:59 (16:42) |
| MPRAGE-$T_1$w/$T_2$w | MPRAGE $T_1$w<br>SPACE $T_2$w | 4:05 + 4:04 (8:09) |
| SPACE-$T_1$w/$T_2$w | SPACE $T_1$w<br>SPACE $T_2$w | 4:11 + 4:04 (8:15) |
| $R_2$ | MESE<br>MPRAGE $T_1$w | 3:22 + 4:05 (7:27) |
| $R_2$' | MEGRE<br>MESE<br>MPRAGE $T_1$w | 5:37 + 3:22 + 4:05 (13:04) |
| $R_2$* | MEGRE<br>MPRAGE $T_1$w | 5:37 + 4:05 (9:42) |
| *q*-Ratio | MEGRE<br>MP2RAGE | 5:37 + 7:59 (13:36) |
| *q*-Ratio-$B_1$c | MEGRE<br>MP2RAGE<br>$B_1^+$ field map | 5:37 + 7:59 + 0:21 (13:57) |
| QSM-SEPIA<br>QSM-X-$R_2^*$<br>QSM-X-$R_2$pnet | MEGRE<br>MPRAGE $T_1$w | 5:37 + 4:05 (9:42) |
| QSM-X-$R_2'$ | MEGRE<br>MESE<br>MPRAGE $T_1$w | 5:37 + 3:22 + 4:05 (13:04) |
| *g*-ihMTsat | ihMTRAGE<br>dMRI<br>MP2RAGE<br>$B_1^+$ field map | 8:22 + 12:48 + 7:59 + 0:21 (29:30) |
| *g*-ihMTR | ihMTRAGE<br>dMRI<br>MPRAGE $T_1$w | 8:22 + 12:48 + 4:05 (25:15) |

**Supplementary Table S1 | Derivatives and Durations.** For each metric analyzed in this study, the MRI acquisitions used as source data in our processing workflows and the cumulative acquisition durations of those acquisitions are shown. Durations are based on the per-session acquisition times reported in **Table 1**. For some metrics, the MPRAGE $T_1$w image was used only as an anatomical reference for registration and spatial normalization. MPRAGE acquisition time is not included for derivatives that already require MP2RAGE because the $T_1$w image generated by MP2RAGE could instead serve as the anatomical reference.

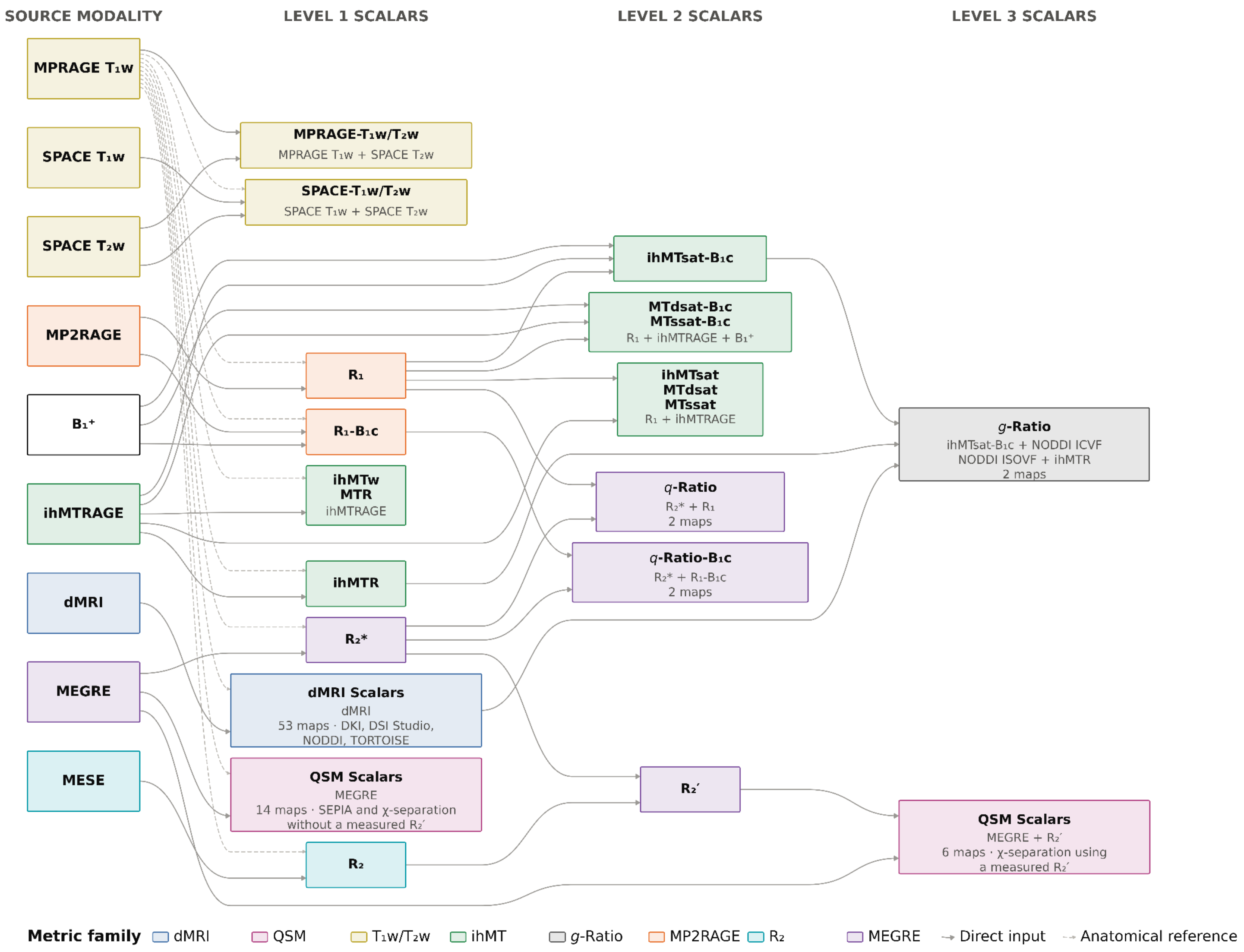


**Supplementary Figure S1 | Derivatives and Source Modalities.** Columns indicate successive levels of derivation, from measures calculated directly from the source acquisitions to higher-order measures that combine previously derived quantities. Arrows indicate dependencies between acquisitions and derived measures; lighter anatomical-reference connections indicate use of anatomical images for registration or spatial normalization rather than direct contribution to the scalar calculation.

| Bundle | Laterality |
|---|---|
| **Association** 36 bundles | |
| Arcuate fasciculus | Bilateral |
| Cingulum frontal parahippocampal | Bilateral |
| Cingulum frontal parietal | Bilateral |
| Cingulum parahippocampal–parietal | Bilateral |
| Cingulum parahippocampal | Bilateral |
| Cingulum parolfactory | Bilateral |
| Extreme capsule | Bilateral |
| Frontal aslant tract | Bilateral |
| Hippocampus alveus | Bilateral |
| Inferior fronto-occipital fasciculus | Bilateral |
| Inferior longitudinal fasciculus | Bilateral |
| Middle longitudinal fasciculus | Bilateral |
| Parietal aslant tract | Bilateral |
| Superior longitudinal fasciculus 1 | Bilateral |
| Superior longitudinal fasciculus 2 | Bilateral |
| Superior longitudinal fasciculus 3 | Bilateral |
| Uncinate fasciculus | Bilateral |
| Vertical occipital fasciculus | Bilateral |
| **Projection—Basal Ganglia** 26 bundles | |
| Acoustic radiation | Bilateral |
| Corticostriatal tract anterior | Bilateral |
| Corticostriatal tract posterior | Bilateral |
| Corticostriatal tract superior | Bilateral |
| Thalamic radiation anterior | Bilateral |

| | |
|---|---|
| Thalamic radiation posterior | Bilateral |
| Thalamic radiation superior | Bilateral |
| Optic radiation | Bilateral |
| Fornix | Bilateral |
| Ansa lenticularis | Bilateral |
| Ansa subthalamica | Bilateral |
| Fasciculus lenticularis | Bilateral |
| Fasciculus subthalamicus | Bilateral |
| **Projection—Brainstem** 20 bundles | |
| Corticospinal tract | Bilateral |
| Corticobulbar tract | Bilateral |
| Corticopontine tract frontal | Bilateral |
| Corticopontine tract parietal | Bilateral |
| Corticopontine tract occipital | Bilateral |
| Medial lemniscus | Bilateral |
| Dentato-rubro-thalamic tract[†] | Decussating bilateral (L→R and R→L) |
| Non-decussating dentato-rubro-thalamic tract | Bilateral |
| Reticular tract | Bilateral |
| Medial forebrain bundle | Bilateral |
| **Commissural** 7 bundles | |
| Anterior commissure frontal[†] | Midline / unpaired |
| Anterior commissure temporal[†] | Midline / unpaired |
| Anterior commissure occipital[†] | Midline / unpaired |
| Corpus callosum forceps minor | Midline / unpaired |
| Corpus callosum body | Midline / unpaired |
| Corpus callosum tapetum | Midline / unpaired |

| | |
|---|---|
| Corpus callosum forceps major | Midline / unpaired |
| **Cerebellar** 7 bundles | |
| Cerebellum | Bilateral |
| Inferior cerebellar peduncle | Bilateral |
| Middle cerebellar peduncle | Midline / unpaired |
| Superior cerebellar peduncle | Midline / unpaired |
| Vermis | Midline / unpaired |

**Supplementary Table S2 | White matter bundles reconstructed by DSI Studio AutoTrack**. Bundles are grouped by anatomical category, with laterality indicated for each tract type. Bilateral tract types yielded separate left- and right-hemisphere bundles, whereas midline tracts were represented by a single bundle. The full reconstruction set comprised 96 bundles. Bundles marked with a dagger (†)—the two crossed dentato-rubro-thalamic tracts and the three anterior commissure bundles—were not reconstructed consistently across all included participant-sessions and were excluded from downstream analyses, leaving 91 white matter bundles.

#### Supplementary Note S1 | *g*-ratio scaling factor calculation

##### Variable definitions

| Symbol | Definition |
|---|---|
| $g$ | Aggregate MRI *g*-ratio |
| FVF | Fiber volume fraction |
| MVF | Unscaled myelin volume fraction proxy |
| $MVF_s$ | Scaled myelin volume fraction |
| $s$ | Multiplicative scaling factor applied to MVF |
| ICVF | Intracellular volume fraction from NODDI |
| ISOVF | Isotropic volume fraction from NODDI |

The aggregate MRI *g*-ratio was defined from the fiber volume fraction (FVF) and the scaled myelin volume fraction ($MVF_s$) as

$$g = \sqrt{\frac{FVF}{FVF + MVF_s}}$$

The fiber volume fraction was estimated from the intracellular volume fraction (ICVF) and isotropic volume fraction (ISOVF), while accounting for the fraction of tissue occupied by myelin:

$$FVF = (1 - MVF_s)(1 - ISOVF)\, ICVF$$

$$MVF_s = s\, MVF$$

##### Derivation

Substituting $MVF_s = s \cdot MVF$ into the FVF expression gives

$$FVF = (1 - s\, MVF)(1 - ISOVF)\, ICVF$$

Substituting this expression into the *g*-ratio equation and squaring both sides yields

$$g^2 = \frac{(1 - s\, MVF)(1 - ISOVF)\, ICVF}{(1 - s\, MVF)(1 - ISOVF)\, ICVF + s\, MVF}$$

Multiplying both sides by the denominator:

$$g^2[(1 - s\, MVF)(1 - ISOVF)\, ICVF + s\, MVF] = (1 - s\, MVF)(1 - ISOVF)\, ICVF$$

Rearranging to isolate the terms containing $s$:

$$g^2 s\, MVF = (1 - g^2)(1 - ISOVF)\, ICVF(1 - s\, MVF)$$

$$g^2 s\, MVF = (1 - g^2)(1 - ISOVF)\, ICVF - (1 - g^2)(1 - ISOVF)\, ICVF\, s\, MVF$$

Collecting the s terms on the left-hand side gives

$$s\, MVF[g^2 + (1 - g^2)(1 - ISOVF)\, ICVF] = (1 - g^2)(1 - ISOVF)\, ICVF$$

Therefore, the scaling factor is

$$s = \frac{(1 - g^2)(1 - ISOVF)\, ICVF}{MVF[g^2 + (1 - g^2)(1 - ISOVF)\, ICVF]}$$

##### Calibration to *g* = 0.7

For a target aggregate *g*-ratio of 0.7, $g^2 = 0.49$ and $1 - g^2 = 0.51$. The expression therefore simplifies to

$$s = \frac{0.51(1 - ISOVF)\, ICVF}{MVF[0.49 + 0.51(1 - ISOVF)\, ICVF]}$$

A numerical value for *s* cannot be obtained from the target *g*-ratio alone; the expression must also be evaluated using the MVF, ICVF, and ISOVF values chosen for calibration. If a single global scaling factor is used, these should be the corresponding summary values used in the analysis. If the expression is evaluated voxelwise, the resulting *s* will vary spatially.

#### Compact implementation form

Defining A = (1 − ISOVF)ICVF, the same result can be written more compactly as

$$A = (1 - ISOVF)\, ICVF$$

$$s = \frac{(1 - g^2)A}{MVF[g^2 + (1 - g^2)A]}$$

For *g* = 0.7:

$$s = \frac{0.51A}{MVF(0.49+0.51A)}$$

| Metric | Description |
| --- | --- |
| ***dMRI*** **53 metrics** | |
| **DKI Micro AD** | Axial diffusivity estimated by DIPY's DKI microstructural model. |
| **DKI Micro ADE** | Axial diffusivity of the extra-cellular compartment estimated by DIPY's DKI microstructural model. |
| **DKI Micro AWF**† | Axonal water fraction estimated by DIPY's DKI microstructural model. |
| **DKI Micro AxonalD** | Axonal diffusivity estimated by DIPY's DKI microstructural model. |
| **DKI Micro KFA** | Kurtosis fractional anisotropy estimated by DIPY's DKI microstructural model. |
| **DKI Micro MD** | Mean diffusivity estimated by DIPY's DKI microstructural model. |
| **DKI Micro RD** | Radial diffusivity estimated by DIPY's DKI microstructural model. |
| **DKI Micro RDE** | Radial diffusivity of the extra-cellular compartment estimated by DIPY's DKI microstructural model. |
| **DKI Micro Tortuosity** | Tortuosity estimated by DIPY's DKI microstructural model. |
| **DKI AD** | Axial diffusivity estimated by DIPY's DKI model. |
| **DKI AK** | Axial kurtosis estimated by DIPY's DKI model. |
| **DKI KFA** | Kurtosis fractional anisotropy estimated by DIPY's DKI model. |
| **DKI MD** | Mean diffusivity estimated by DIPY's DKI model. |
| **DKI MK** | Mean kurtosis estimated by DIPY's DKI model. |
| **DKI MKT**† | Mean of the kurtosis tensor estimated by DIPY's DKI model. |
| **DKI RD** | Radial diffusivity estimated by DIPY's DKI model. |
| **DKI RK**† | Radial kurtosis estimated by DIPY's DKI model. |
| **DKI Tensor FA** | Fractional anisotropy estimated by DIPY's DKI model. |
| **DSI Studio GQI GFA**† | Generalized fractional anisotropy estimated by DSI Studio's GQI fit. |
| **DSI Studio GQI ISO** | Isotropic diffusion estimated by DSI Studio's GQI fit. |
| **DSI Studio GQI QA** | Quantitative anisotropy estimated by DSI Studio's GQI fit. |
| **DSI Studio Tensor AD** | Axial diffusivity estimated by DSI Studio's tensor fit. |
| **DSI Studio Tensor FA** | Fractional anisotropy estimated by DSI Studio's tensor fit. |
| **DSI Studio Tensor MD** | Mean diffusivity estimated by DSI Studio's tensor fit. |
| **DSI Studio Tensor RD** | Radial diffusivity estimated by DSI Studio's tensor fit. |
| **NODDI ICVF Modulated** | Intracellular volume fraction modulated by tissue fraction estimated by AMICO's NODDI model. |
| **NODDI ICVF**† | Intracellular volume fraction estimated by AMICO's NODDI model. |
| **NODDI ISOVF** | Isotropic volume fraction estimated by AMICO's NODDI model. |

| | |
|---|---|
| **NODDI OD Modulated** | Orientation dispersion index modulated by tissue fraction estimated by AMICO's NODDI model. |
| **NODDI OD** | Orientation dispersion index estimated by AMICO's NODDI model. |
| **NODDI ICVF Modulated (GM)** | Intracellular volume fraction modulated by tissue fraction estimated by AMICO's gray matter NODDI model with gray matter diffusivity constants. |
| **NODDI ICVF (GM)†** | Intracellular volume fraction estimated by AMICO's NODDI model with gray matter diffusivity constants. |
| **NODDI ISOVF (GM)** | Isotropic volume fraction estimated by AMICO's gray matter NODDI model with gray matter diffusivity constants. |
| **NODDI OD Modulated (GM)** | Orientation dispersion index modulated by tissue fraction estimated by AMICO's gray matter NODDI model with gray matter diffusivity constants. |
| **NODDI OD (GM)** | Orientation dispersion index estimated by AMICO's gray matter NODDI model with gray matter diffusivity constants. |
| **TORTOISE MAP-MRI NG†** | Non-Gaussianity estimated by TORTOISE's MAP-MRI model. |
| **TORTOISE MAP-MRI NG \|\|** | Parallel non-Gaussianity estimated by TORTOISE's MAP-MRI model. |
| **TORTOISE MAP-MRI NG ⊥†** | Perpendicular non-Gaussianity estimated by TORTOISE's MAP-MRI model. |
| **TORTOISE MAP-MRI PA** | Propagator anisotropy estimated by TORTOISE's MAP-MRI model. |
| **TORTOISE MAP-MRI PAth** | Angular measure of propagator anisotropy derived from TORTOISE's MAP-MRI model, prior to the nonlinear transformation used to calculate propagator anisotropy (PA). |
| **TORTOISE MAP-MRI RTAP†** | Return to axis probability estimated by TORTOISE's MAP-MRI model. |
| **TORTOISE MAP-MRI RTOP†** | Return to origin probability estimated by TORTOISE's MAP-MRI model. |
| **TORTOISE MAP-MRI RTPP** | Return to plane probability estimated by TORTOISE's MAP-MRI model. |
| **TORTOISE MAP-MRI AD** | Axial diffusivity estimated by TORTOISE's tensor model from inner shells only. |
| **TORTOISE MAP-MRI FA†** | Fractional anisotropy estimated by TORTOISE's tensor model from inner shells only. |
| **TORTOISE MAP-MRI LI** | LI estimated by TORTOISE's tensor model from inner shells only. |
| **TORTOISE MAP-MRI RD†** | Radial diffusivity estimated by TORTOISE's tensor model from inner shells only. |
| **TORTOISE MAP-MRI MD†** | Mean diffusivity estimated by TORTOISE's tensor model from inner shells only. |
| **TORTOISE Tensor AD** | Axial diffusivity estimated by TORTOISE's tensor model from all shells. |
| **TORTOISE Tensor FA** | Fractional anisotropy estimated by TORTOISE's tensor model from all shells. |
| **TORTOISE Tensor LI** | LI estimated by TORTOISE's tensor model from all shells. |
| **TORTOISE Tensor RD** | Radial diffusivity estimated by TORTOISE's tensor model from all shells. |
| **TORTOISE Tensor MD** | Mean diffusivity estimated by TORTOISE's tensor model from all shells. |
| ***ihMT*** **5 metrics** | |
| **ihMTw** | Inhomogeneous magnetization transfer-weighted map estimated from ihMTRAGE. |
| **ihMTR†** | Inhomogeneous magnetization transfer ratio map estimated from ihMTRAGE. |
| **MTR** | Magnetization transfer ratio map estimated from ihMTRAGE (single saturation data). |
| **ihMTsat** | Inhomogeneous magnetization transfer saturation map estimated from ihMTRAGE. |

| | |
|---|---|
| **ihMTsat-$B_1c$†** | Inhomogeneous magnetization transfer saturation map normalized by squared $B_1^+$ estimated from ihMTRAGE and $B_1^+$ field map. |
| **$R_2$†** 1 metric | Irreversible transverse relaxation rate, in $seconds^{-1}$. |
| ***MESE/MEGRE*** 2 metrics | |
| **$R_2$'-E5** | Reversible transverse relaxation rate, in $seconds^{-1}$, estimated from all five echoes of the MEGRE scan and the MESE scan. |
| **$R_2$'-E4** | Reversible transverse relaxation rate, in $seconds^{-1}$, estimated from the last four echoes of the MEGRE scan and the MESE scan. |
| ***MP2RAGE*** 2 metrics | |
| **$R_1$†** | $R_1$ map estimated from MP2RAGE, without $B_1$-correction. In $seconds^{-1}$. |
| **$R_1$-$B_1c$†** | $R_1$ map estimated from MP2RAGE, with $B_1$-correction. In $seconds^{-1}$. |
| ***$T_1w/T_2w$*** 2 metrics | |
| **MPRAGE-$T_1w/T_2w$†** | $T_1w/T_2w$ ratio map estimated from MPRAGE $T_1w$ and SPACE $T_2w$. |
| **SPACE-$T_1w/T_2w$†** | $T_1w/T_2w$ ratio map estimated from SPACE $T_1w$ and SPACE $T_2w$. |
| ***g-Ratio*** 2 metrics | |
| ***g*-ihMTsat†** | *g*-ratio map estimated from ihMTsat-$B_1c$, ISOVF, and ICVF maps. |
| ***g*-ihMTR†** | *g*-ratio map estimated from ihMTR, ISOVF, and ICVF maps. |
| ***MEGRE*** 6 metrics | |
| **$R_2$*-E5†** | Apparent transverse relaxation rate, in $seconds^{-1}$, estimated from all five echoes of the MEGRE scan. |
| **$R_2$*-E4** | Apparent transverse relaxation rate, in $seconds^{-1}$, estimated from the last four echoes of the MEGRE scan. |
| ***q*-Ratio-E5** | $R_1$ x $R_2$*, estimated from the uncorrected $R_1$ derived from the MP2RAGE scan and the $R_2$* derived from all five echoes of the MEGRE scan. In $seconds^{-2}$. |
| ***q*-Ratio-E5-$B_1c$†** | $R_1$ x $R_2$*, estimated from the $B_1$-corrected $R_1$ derived from the MP2RAGE scan and the $R_2$* derived from all five echoes of the MEGRE scan. In $seconds^{-2}$. |
| ***q*-Ratio-E4** | $R_1$ x $R_2$*, estimated from the uncorrected $R_1$ derived from the MP2RAGE scan and the $R_2$* derived from the last four echoes of the MEGRE scan. In $seconds^{-2}$. |
| ***q*-Ratio-E4-$B_1c$** | $R_1$ x $R_2$*, estimated from the $B_1$-corrected $R_1$ derived from the MP2RAGE scan and the $R_2$* derived from the last four echoes of the MEGRE scan. In $seconds^{-2}$. |
| ***QSM*** 20 metrics | |
| **QSM-SEPIA-E5-χ†** | Chi map from SEPIA toolbox, using all five echoes of the MEGRE scan. |
| **QSM-SEPIA-E4-χ** | Chi map from SEPIA toolbox, using the last four echoes of the MEGRE scan. |
| **QSM-χ-$R_2$p-E5-χ†** | Chi map from χ-sep-net toolbox, using all five echoes of the MEGRE scan and $R_2$’ map estimated from MEGRE and MESE scans. |
| **QSM-χ-$R_2$p-E4-χ** | Chi map from χ-sep-net toolbox, using the last four echoes of the MEGRE scan and $R_2$’ map estimated from MEGRE and MESE scans. |
| **QSM-χ-$R_2$p-E5-Para†** | Paramagnetic susceptibility map from χ-sep-net toolbox, using all five echoes of the MEGRE scan and $R_2$’ map estimated from MEGRE and MESE scans. |
| **QSM-χ-$R_2$p-E4-Para** | Paramagnetic susceptibility map from χ-sep-net toolbox, using the last four echoes of the MEGRE scan and $R_2$’ map estimated from MEGRE and MESE scans. |

| | |
|---|---|
| **QSM-χ-$R_2$p-E5-Dia†** | Diamagnetic susceptibility map from χ-sep-net toolbox, using all five echoes of the MEGRE scan and $R_2$' map estimated from MEGRE and MESE scans. |
| **QSM-χ-$R_2$p-E4-Dia** | Diamagnetic susceptibility map from χ-sep-net toolbox, using the last four echoes of the MEGRE scan and $R_2$' map estimated from MEGRE and MESE scans. |
| **QSM-χ-$R_2$pnet-E5-χ** | Chi map from χ-sep-net toolbox, using all five echoes of the MEGRE scan and $R_2$pnet algorithm. |
| **QSM-χ-$R_2$pnet-E4-χ** | Chi map from χ-sep-net toolbox, using the last four echoes of the MEGRE scan and $R_2$pnet algorithm. |
| **QSM-χ-$R_2$pnet-E5-Para** | Paramagnetic susceptibility map from χ-sep-net toolbox, using all five echoes of the MEGRE scan and $R_2$pnet algorithm. |
| **QSM-χ-$R_2$pnet-E4-Para** | Paramagnetic susceptibility map from χ-sep-net toolbox, using the last four echoes of the MEGRE scan and $R_2$pnet algorithm. |
| **QSM-χ-$R_2$pnet-E5-Dia** | Diamagnetic susceptibility map from χ-sep-net toolbox, using all five echoes of the MEGRE scan and $R_2$pnet algorithm. |
| **QSM-χ-$R_2$pnet-E4-Dia** | Diamagnetic susceptibility map from χ-sep-net toolbox, using the last four echoes of the MEGRE scan and $R_2$pnet algorithm. |
| **QSM-χ-$R_2$*-E5-χ** | Chi map from χ-sep-net toolbox, using all five echoes of the MEGRE scan and $R_2$* map estimated from MEGRE scan. |
| **QSM-χ-$R_2$*-E4-χ** | Chi map from χ-sep-net toolbox, using the last four echoes of the MEGRE scan and $R_2$* map estimated from MEGRE scan. |
| **QSM-χ-$R_2$*-E5-Para** | Paramagnetic susceptibility map from χ-sep-net toolbox, using all five echoes of the MEGRE scan and $R_2$* map estimated from MEGRE scan. |
| **QSM-χ-$R_2$*-E4-Para** | Paramagnetic susceptibility map from χ-sep-net toolbox, using the last four echoes of the MEGRE scan and $R_2$* map estimated from MEGRE scan. |
| **QSM-χ-$R_2$*-E5-Dia** | Diamagnetic susceptibility map from χ-sep-net toolbox, using all five echoes of the MEGRE scan and $R_2$* map estimated from MEGRE scan. |
| **QSM-χ-$R_2$*-E4-Dia** | Diamagnetic susceptibility map from χ-sep-net toolbox, using the last four echoes of the MEGRE scan and $R_2$* map estimated from MEGRE scan. |

**Supplementary Table S3 | Complete set of metrics analyzed in the supplementary materials.** A total of 93 scalar metrics spanning all processing workflows were evaluated in the supplementary analyses. These comprise most scalar outputs generated by the processing workflows. Selected outputs, including individual diffusion-tensor components and $b_0$ signal amplitude, were excluded because they were not considered relevant to the study of myelin-sensitive imaging. Metrics are grouped according to their acquisition or modeling family. NODDI metrics labeled “GM” were obtained using the gray matter-optimized NODDI fit. Metrics marked with a dagger (†) were included in the 27-metric set used for the primary analyses.

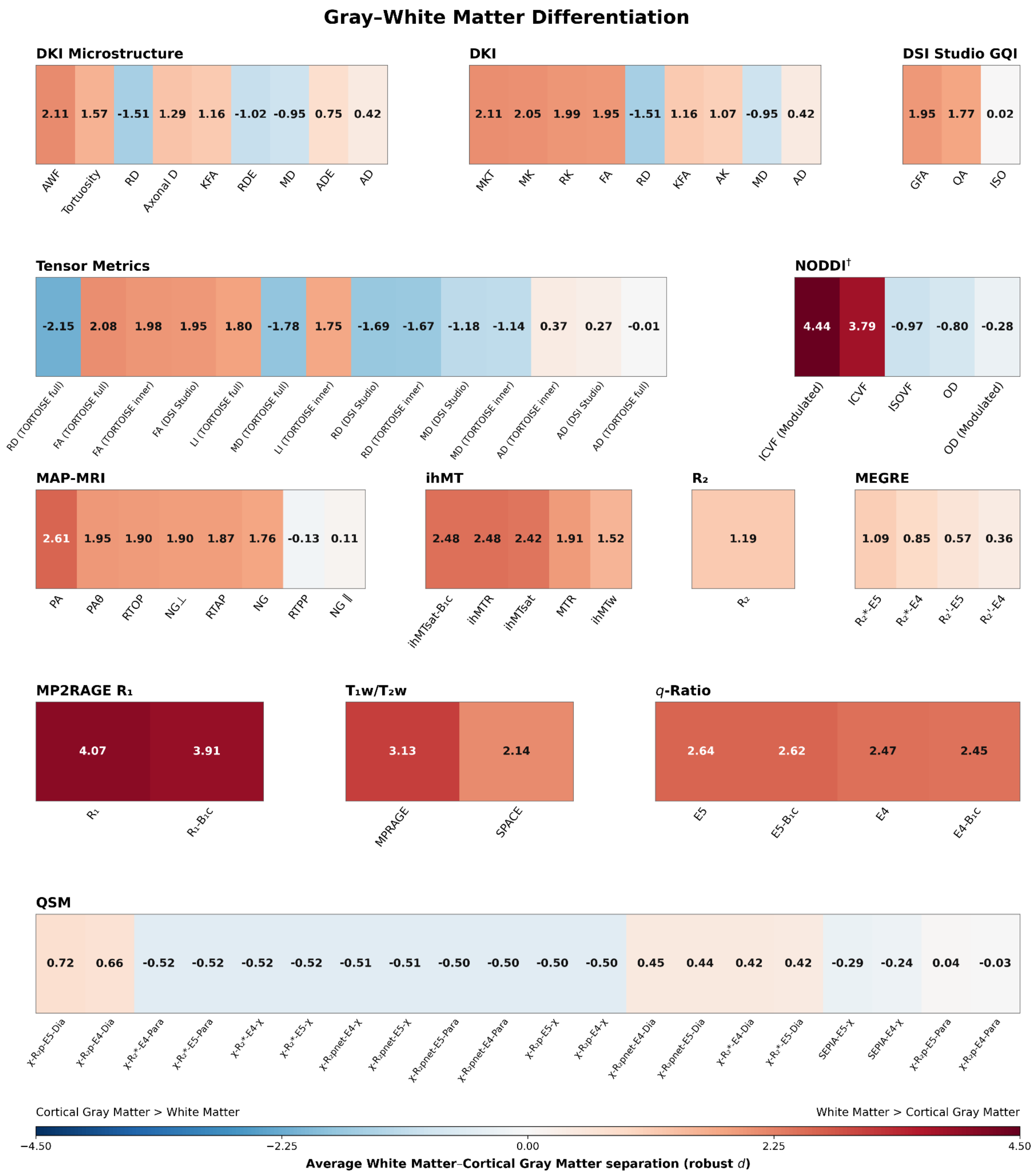


**Supplementary Figure S2 | Gray–white matter differentiation across the expanded metric set.** Gray–white matter differentiation was quantified voxelwise within each participant and session using a robust standardized effect size, calculated as the difference between white matter and cortical gray matter medians divided by a pooled median-absolute-deviation-based estimate of dispersion. Finite, nonzero values were retained after excluding extreme values using an absolute robust *z* score threshold of 6. Session-level effect sizes were averaged within participants and subsequently summarized across participants. Positive values indicate higher metric values in white matter than cortical gray matter, whereas negative values indicate higher values in cortical gray matter. The dagger (†) indicates that the NODDI comparison used two different tissue-specific model fits.

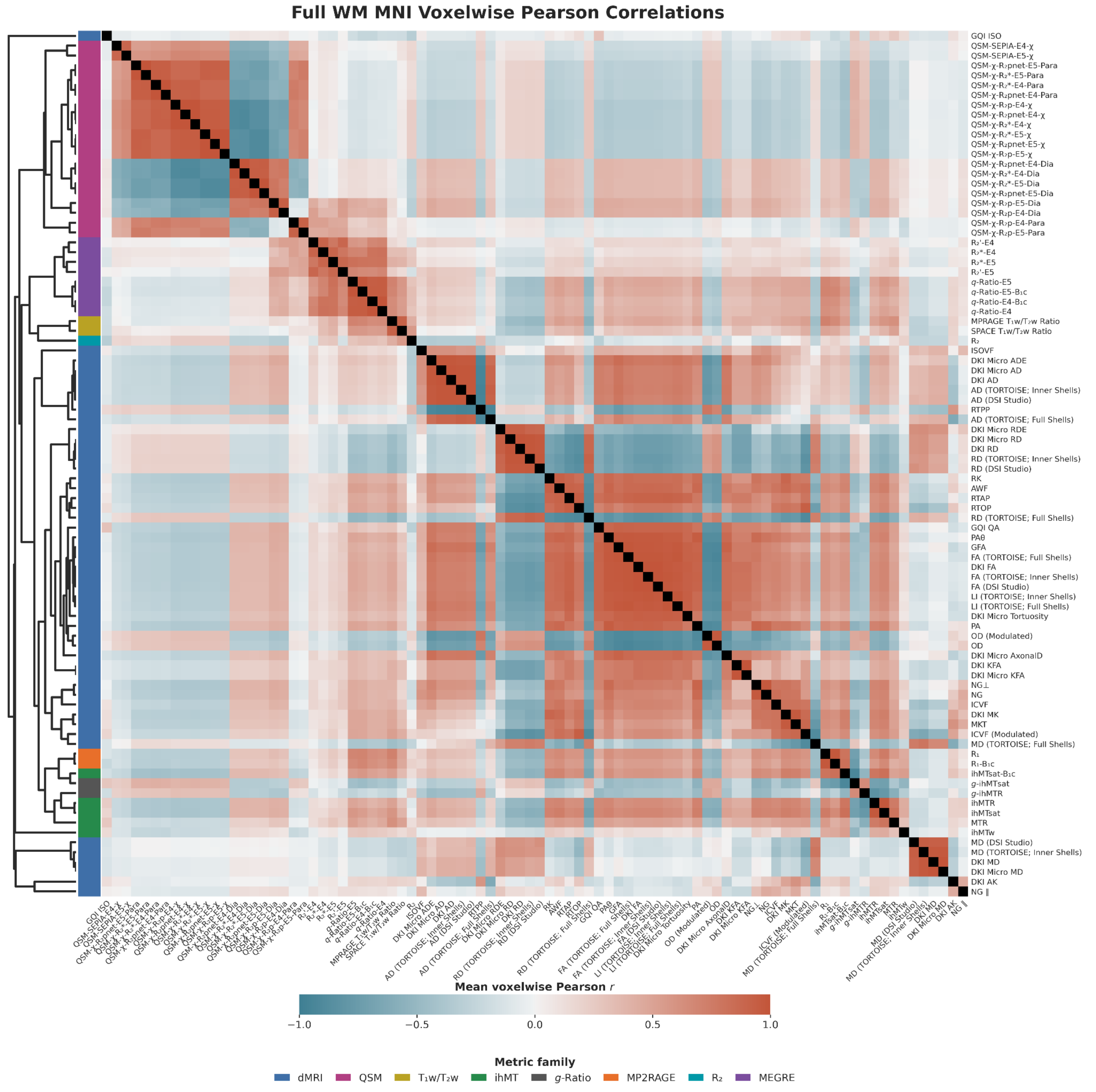


**Supplementary Figure S3 | Voxelwise intermetric spatial correlations in white matter across the expanded metric set.** Pearson correlations were calculated across pairwise-valid white matter voxels separately within each participant and session. Pairwise correlation coefficients were Fisher *z*-transformed and averaged across available sessions within participants and then across participants, giving each participant equal weight, before transformation back to Pearson correlation coefficients. The matrix shows the resulting group-average correlations for all eligible metrics in the expanded supplementary metric set. The dendrogram shows hierarchical organization of the displayed correlation profiles, and colored annotations denote metric families.

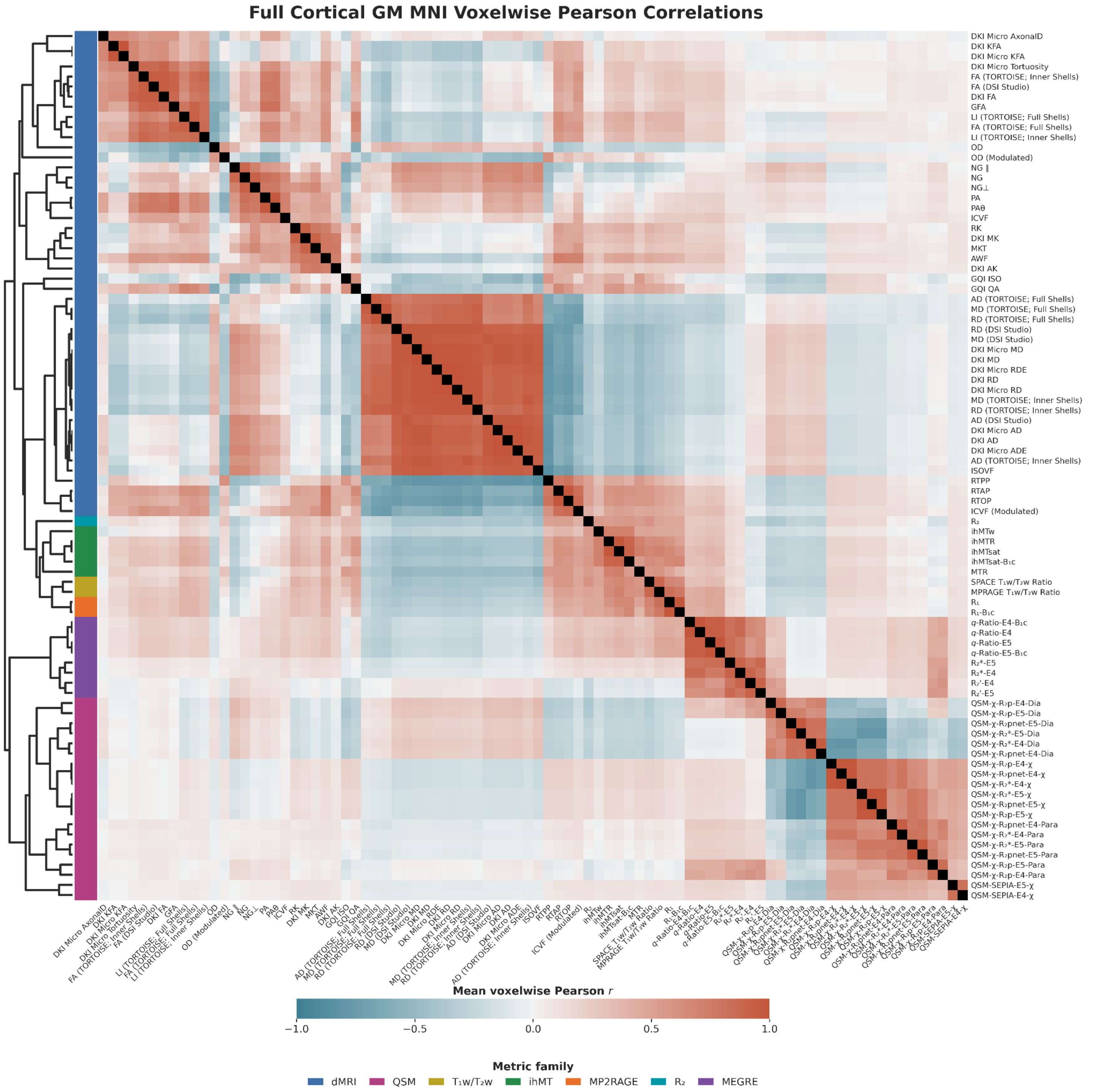


**Supplementary Figure S4 | Voxelwise intermetric spatial correlations in cortical gray matter across the expanded metric set.** Pearson correlations were calculated across pairwise-valid cortical gray matter voxels separately within each participant and session. Pairwise correlation coefficients were Fisher $z$-transformed and averaged across available sessions within participants and then across participants, giving each participant equal weight, before transformation back to Pearson correlation coefficients. The matrix shows the resulting group-average correlations for all eligible metrics in the expanded supplementary metric set. The dendrogram shows hierarchical organization of the displayed correlation profiles, and colored annotations denote metric families.

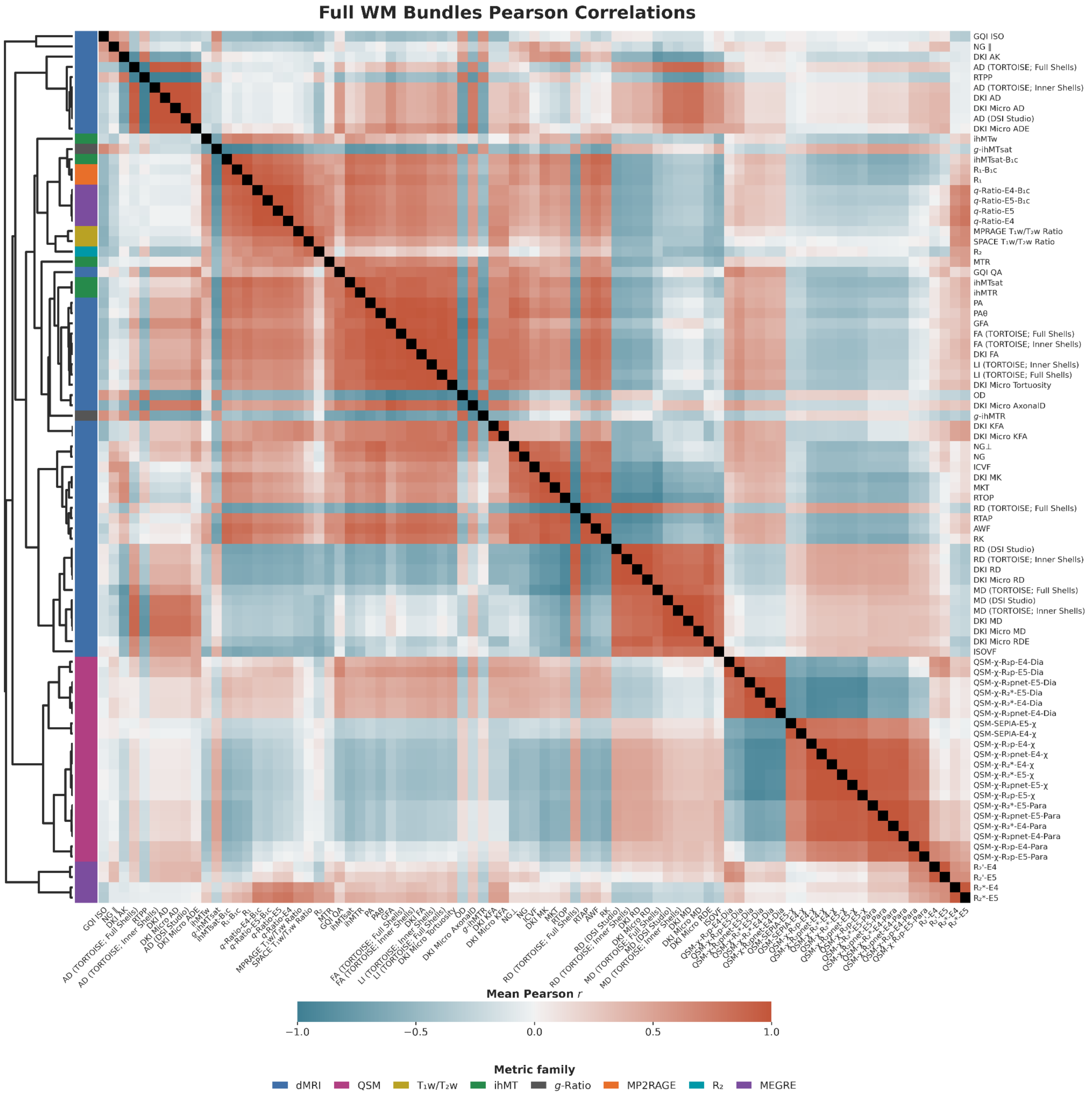


**Supplementary Figure S5 | Bundle-wise intermetric spatial correlations in white matter across the expanded metric set.** Pearson correlations were calculated across pairwise-valid white matter bundles separately within each participant and session. Pairwise correlation coefficients were Fisher *z*-transformed and averaged across available sessions within participants and then across participants, giving each participant equal weight, before transformation back to Pearson correlation coefficients. The matrix shows the resulting group-average correlations for all eligible metrics in the expanded supplementary metric set. The dendrogram shows hierarchical organization of the displayed correlation profiles, and colored annotations denote metric families.

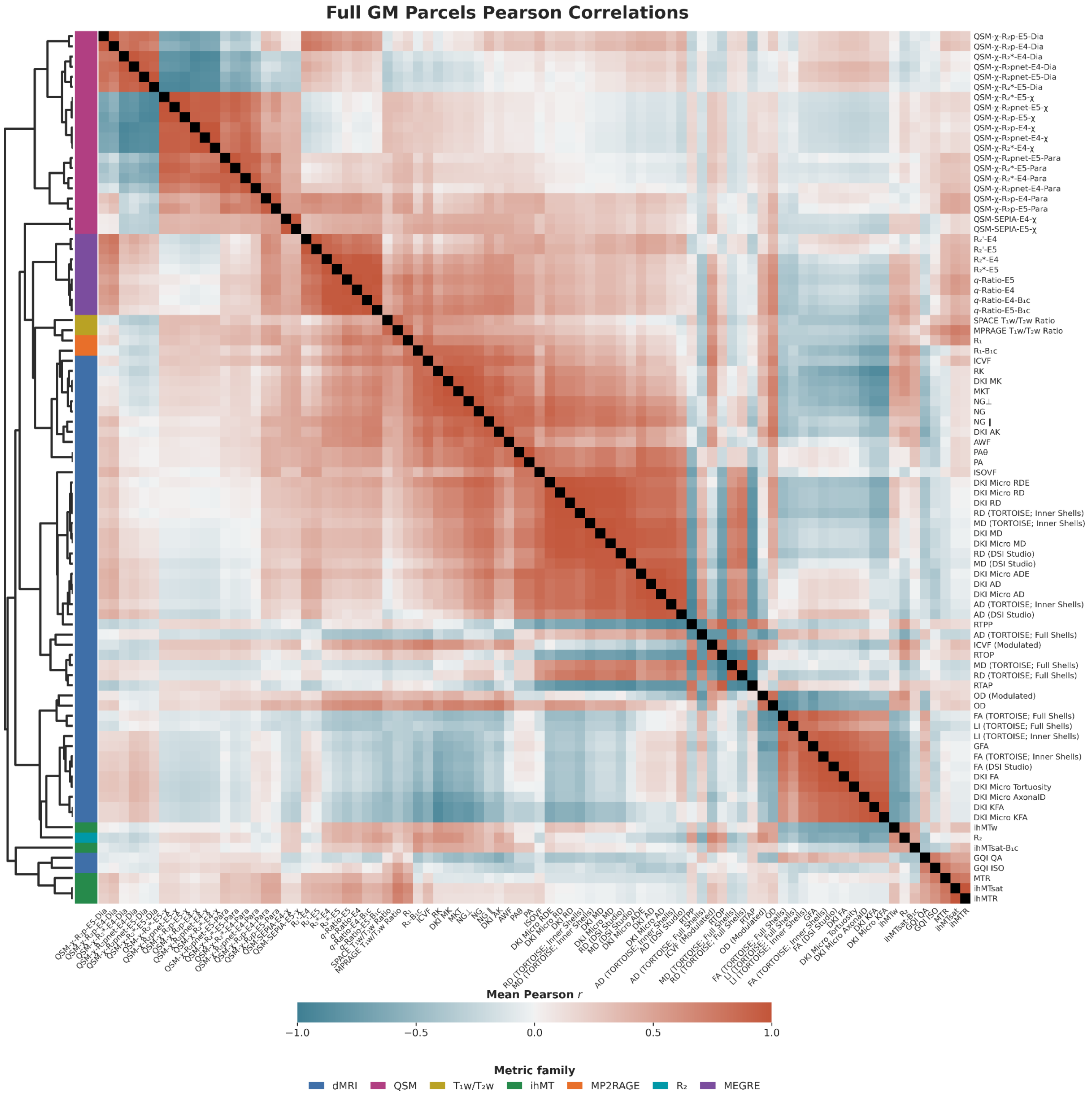


**Supplementary Figure S6 | Parcel-wise intermetric spatial correlations in cortical gray matter across the expanded metric set.** Pearson correlations were calculated across pairwise-valid cortical gray matter parcels separately within each participant and session. Pairwise correlation coefficients were Fisher z-transformed and averaged across available sessions within participants and then across participants, giving each participant equal weight, before transformation back to Pearson correlation coefficients. The matrix shows the resulting group-average correlations for all eligible metrics in the expanded supplementary metric set. The dendrogram shows hierarchical organization of the displayed correlation profiles, and colored annotations denote metric families.

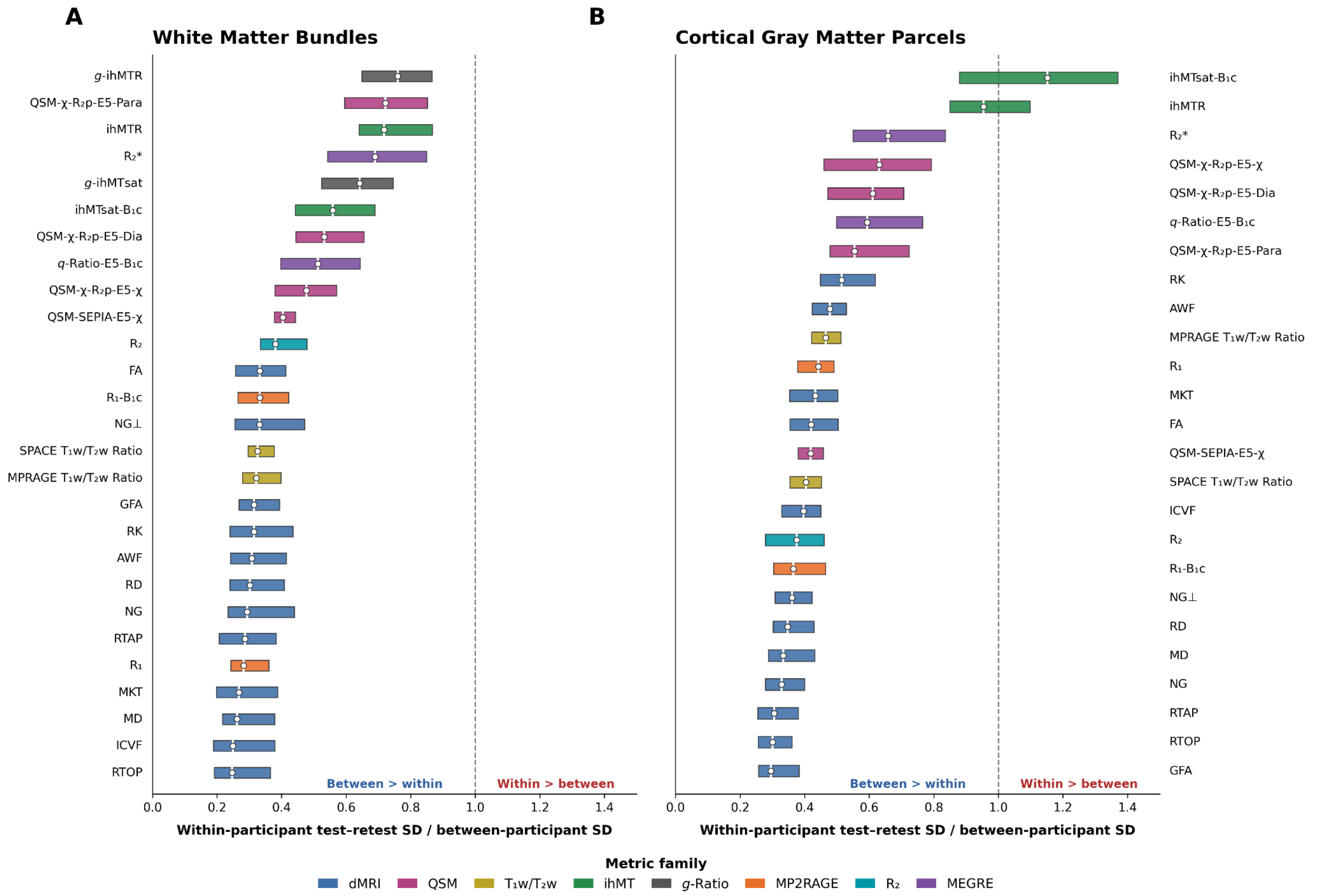


**Supplementary Figure S7 | Within- relative to between-participant variability across regional measurements.** For each metric and region, within-participant test–retest SD was estimated as the square root of the residual mean square from the participant-by-session ANOVA, and between-participant SD was calculated as the SD of participant means across sessions. White circles indicate the median and colored bars span the interquartile range across **(A)** white matter bundles or **(B)** cortical gray matter parcels. The dashed line at 1 indicates equal within- and between-participant variability; values above 1 indicate greater within-participant than between-participant variability. Colors denote metric families.

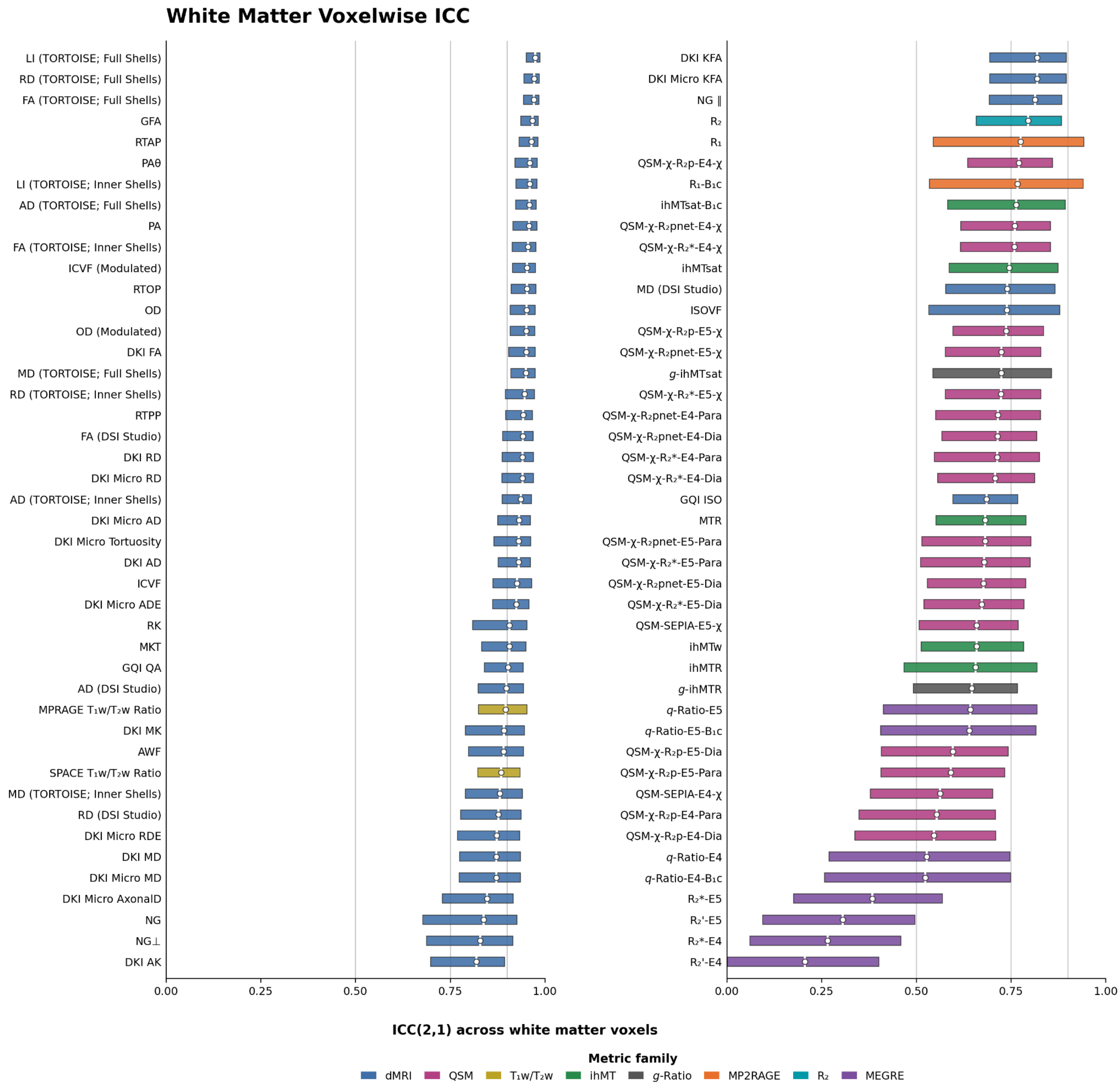


**Supplementary Figure S8 |** Voxelwise test–retest reliability in white matter across the expanded metric set. Test–retest reliability was quantified using ICC(2,1), corresponding to a two-way random-effects model of absolute agreement for a single measurement. ICCs were estimated independently at each voxel within the fixed group-level white matter mask using participants for whom both sessions of the corresponding source acquisition were available and passed quality assurance. For each metric, the white circle indicates the median ICC and the horizontal bar spans the interquartile range of ICC values across white matter voxels. Metrics are ordered by median ICC, and colors denote metric families.

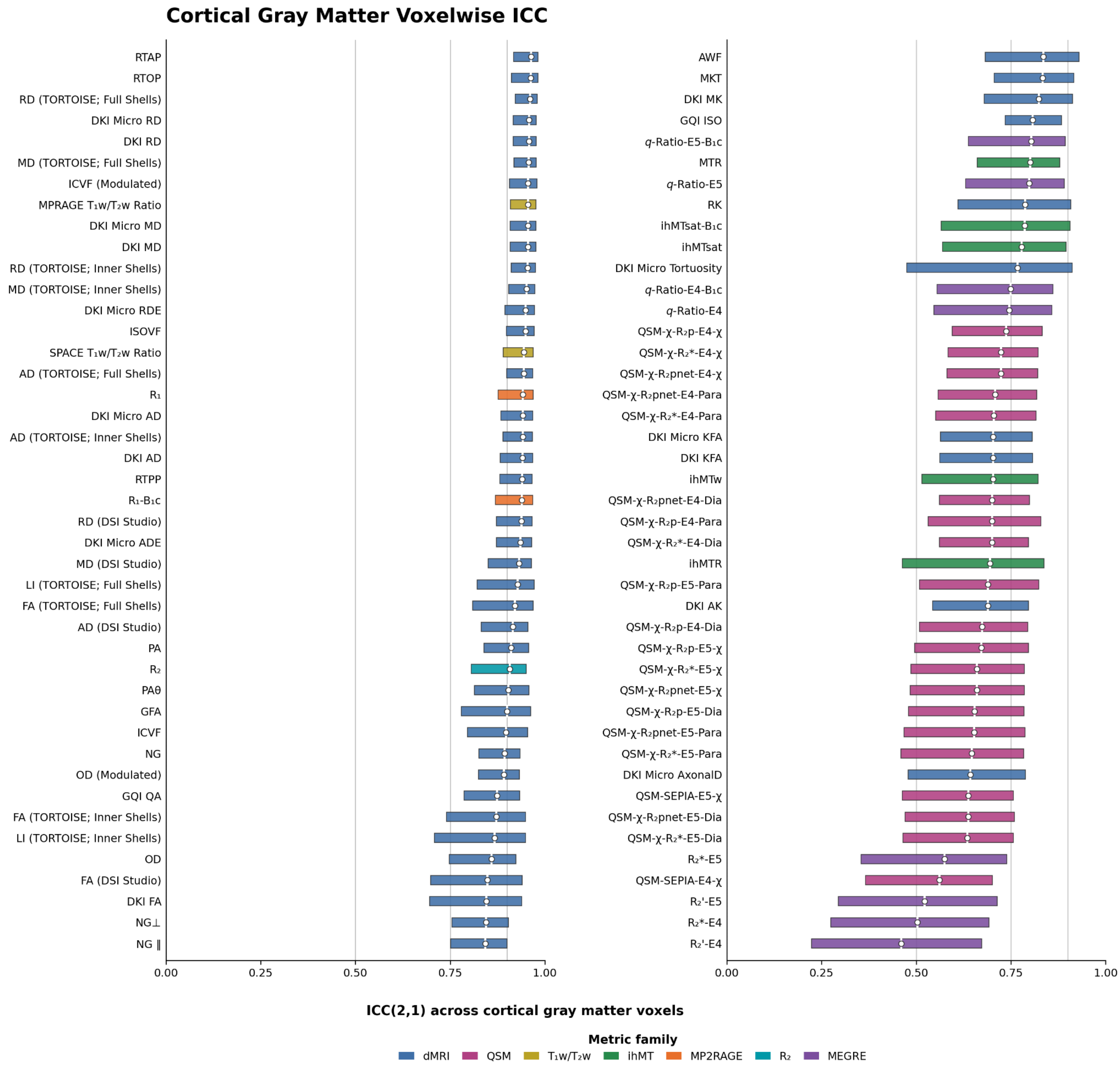


**Supplementary Figure S9 | Voxelwise test–retest reliability in cortical gray matter across the expanded metric set.** Test–retest reliability was quantified using ICC(2,1), corresponding to a two-way random-effects model of absolute agreement for a single measurement. ICCs were estimated independently at each voxel within the fixed group-level cortical gray matter mask using participants for whom both sessions of the corresponding source acquisition were available and passed quality assurance. For each metric, the white circle indicates the median ICC and the horizontal bar spans the interquartile range of ICC values across cortical gray matter voxels. Metrics are ordered by median ICC, and colors denote metric families.

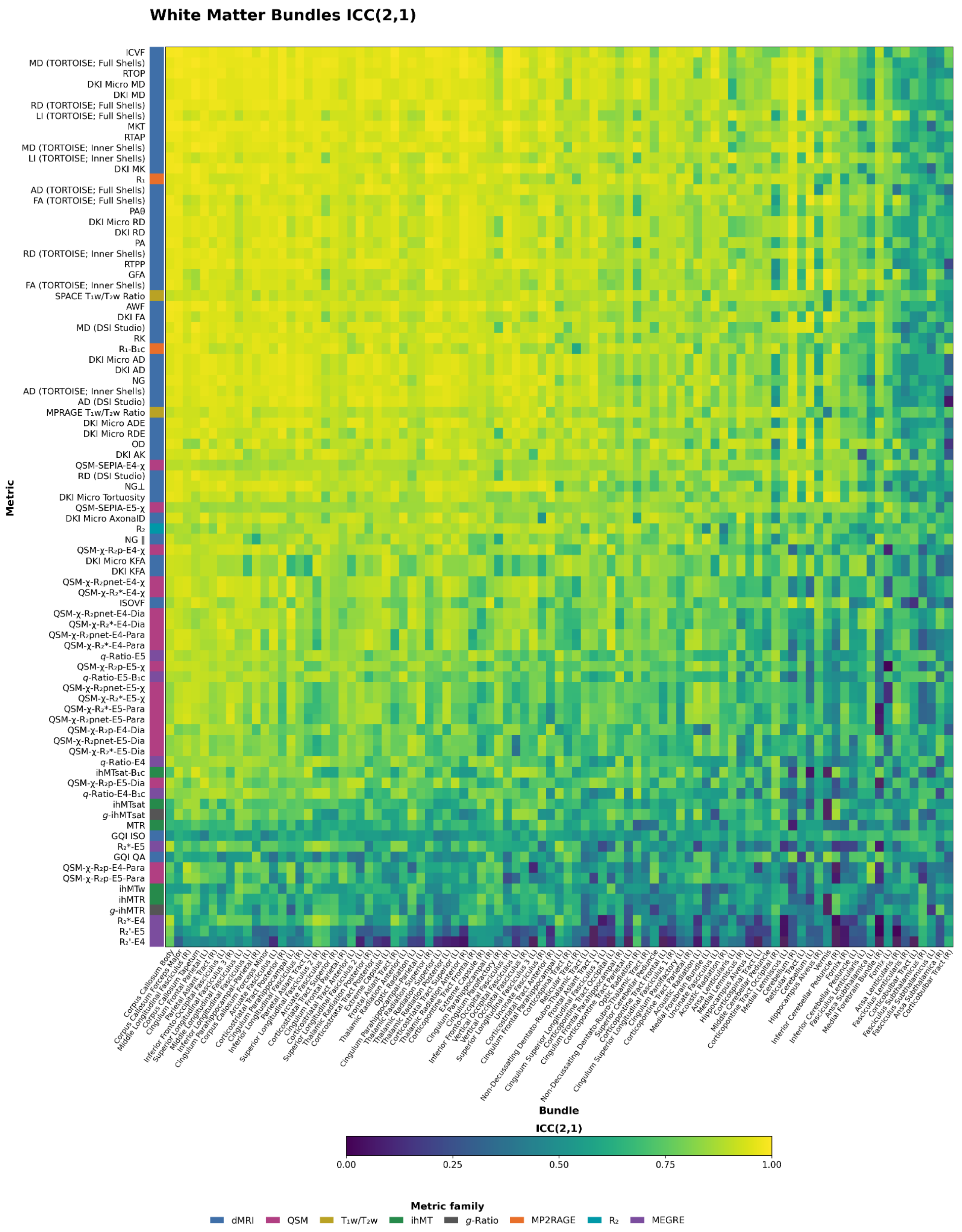


**Supplementary Figure S10 | Bundle-wise test–retest reliability in white matter across the expanded metric set.** Test–retest reliability was quantified using ICC(2,1) independently for each metric and white matter bundle. Each heatmap cell shows the ICC estimated across participants for one metric in one of the 91 participant-specific white matter bundles retained for analysis; only participants with both corresponding sessions available and passing quality assurance contributed to a given metric. Higher values indicate greater test–retest reliability. The annotation beside the matrix denotes the metric family.

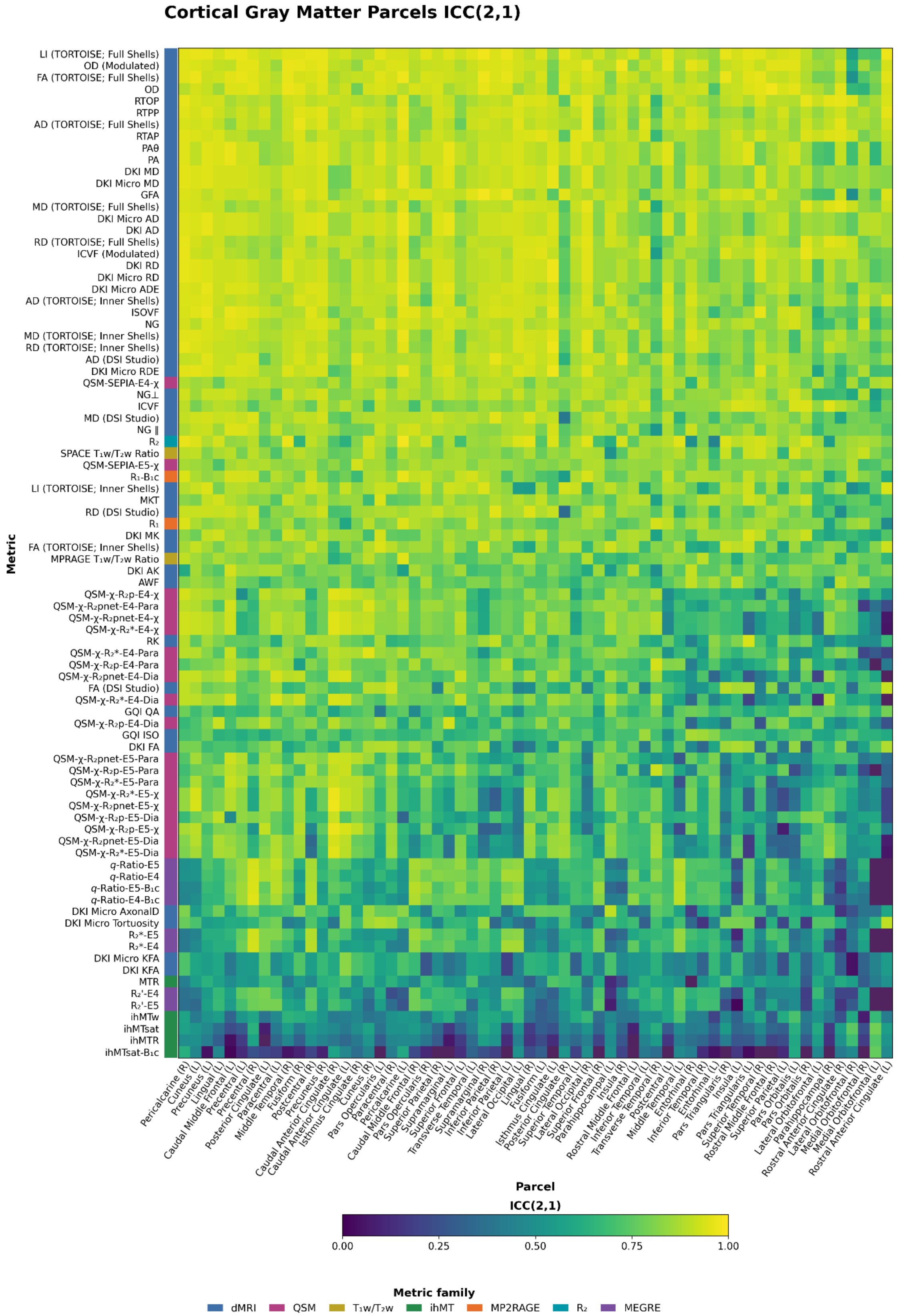


**Supplementary Figure S11 | Parcel-wise test–retest reliability in cortical gray matter across the expanded metric set.** Test–retest reliability was quantified using ICC(2,1) independently for each metric and cortical parcel. Each heatmap cell shows the ICC estimated across participants for one metric in one of the 62 participant-specific Desikan–Killiany–Tourville cortical parcels; only participants with both corresponding sessions available and passing quality assurance contributed to a given metric. Higher values indicate greater test–retest reliability. The annotation beside the matrix denotes the metric family.

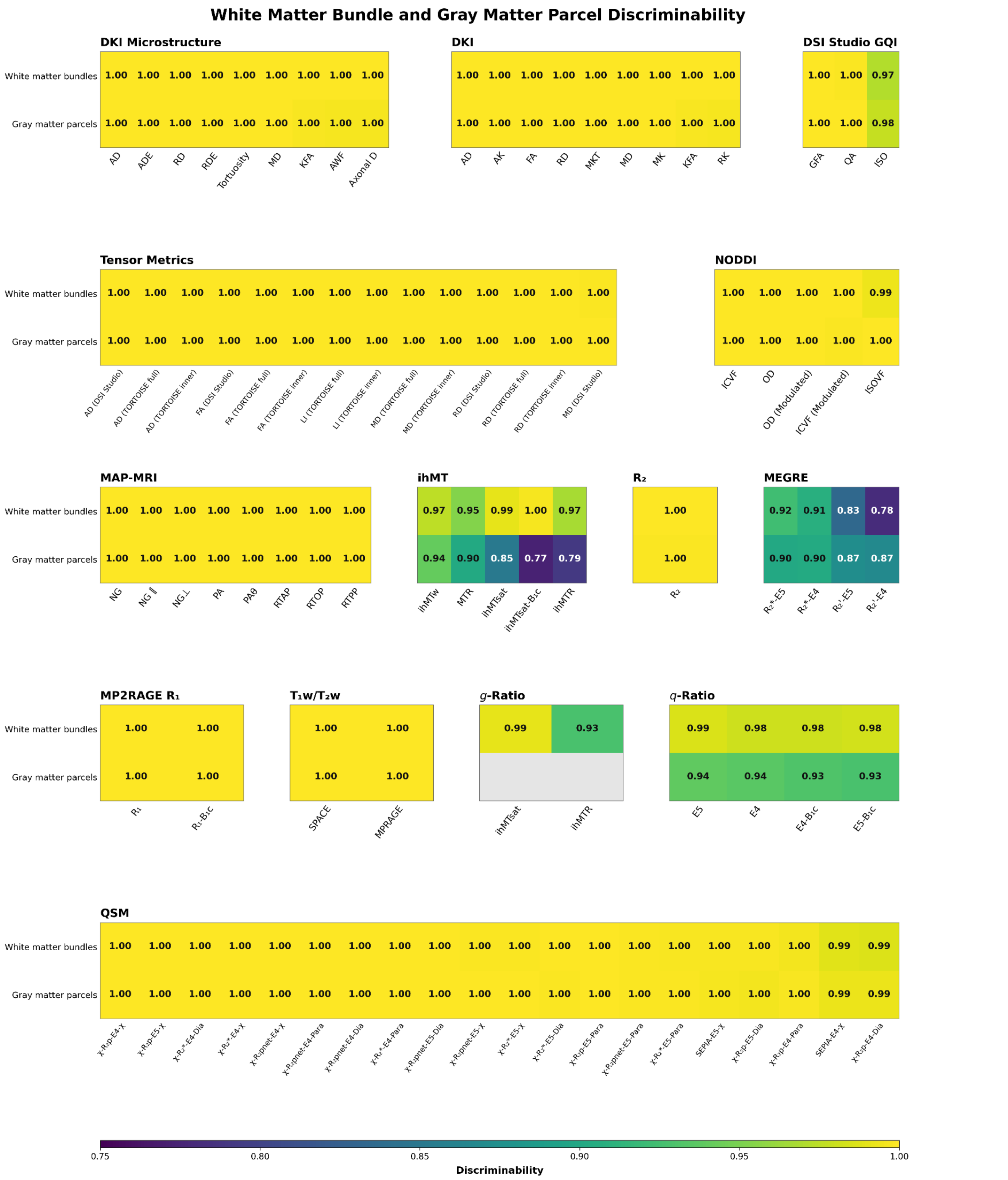


**Supplementary Figure S12 | Discriminability of white matter bundle and cortical gray matter parcel profiles across the expanded metric set.** Cells show discriminability for white matter bundles and cortical gray matter parcels, calculated separately, with values approaching 1 indicating greater participant specificity. Metrics are grouped by processing or modeling family. Gray cells for *g*-ratio measures indicate they were analyzed only in white matter.
